# Quantitative profiling of protease specificity

**DOI:** 10.1101/2020.06.30.179820

**Authors:** B. I. Ratnikov, P. Cieplak, A. G. Remacle, E. Nguyen, J. W. Smith

## Abstract

Proteases comprise an important class of enzymes, whose activity is central to many physiologic and pathologic processes. Detailed knowledge of protease specificity is key to understanding their function. Although many methodologies have been developed to profile specificities of proteases, few have the diversity and quantitative grasp necessary to fully define specificity of a protease, both in terms of substrate numbers and their catalytic efficiencies. We have developed a concept of “selectome”, which defines the set of substrates that uniquely represents specificity of a protease. We applied it to two closely related members of the Matrixin family – MMP-2 and MMP-9 by using substrate phage display coupled with Next Generation Sequencing and information theory-based data analysis. We have also derived a quantitative measure of substrate specificity, which accounts for both the numbers and relative catalytic efficiencies of substrates. Using these advances greatly facilitates uncovering selectivity between closely related members of protease families and provides insight into to the degree of contribution of catalytic cleft specificity to protein substrate recognition, thus providing basis to overcoming two of the major challenges in the field of proteolysis: 1) development of highly selective activity probes and inhibitors for studying proteases with overlapping specificities, and 2) distinguishing targeted proteolysis from bystander proteolytic events.

## Introduction

Proteolytic enzymes a.k.a. proteases or peptidases, whose catalytic function is hydrolysis of peptide bonds, can be classified based on catalytic mechanism into serine, threonine, cysteine, aspartic, glutamic and metalloproteinases [1]. Proteolytic enzymes listed in the MEROPS database are members of 268 gene families and their number is growing as the number of sequenced genomes increases [2]. In humans, proteases comprise approximately 3% of the protein-coding genome and are represented by over 560 unique proteins [3]. Proteases are involved in all aspects of biology from embryonic development to programmed cell death and cellular protein recycling and therefore are an integral part of proteolytic pathways that connect different biological processes into functional networks [3-8]. Protease activity has to be tightly regulated and newly synthesized enzymes often require proenzyme activation and mature proteases are subject to inhibition by a variety of endogenous inhibitors, as deleterious consequences of uncontrolled proteolysis can be devastating [4, 9].

Proteolysis is particularly significant from a systems biology perspective because it is the only irreversible post-translational modification. A proteolytic cleavage is thus a committed step in the function of networks and pathways. Yet, proteases and proteolysis present unique features/characteristics that have made them difficult to study from a systems perspective. These include: 1) redundancy and specificity overlap between proteases belonging to the same families [4, 10], 2) overlapping specificities of proteases belonging to different families and classes [4, 11], 3) difficulty in distinguishing physiologically relevant cleavages from coincidental proteolytic events [3, 4], 4) lack of information about selectivity due to insufficiency of the tools currently available for their study [4, 12-15].

Systems biology approach to studying protease function, which is key to understanding their roles in a broader context of the cell and the whole organism, is facilitated by rapidly accumulating knowledge of protease substrates [2, 3, 16]. At the center of proteolysis is the recognition of substrate at the catalytic cleft. In most cases this site is the primary regulatory point for substrate recognition and selectivity. A full understanding of substrate recognition at this site has the potential to provide information necessary to move the study of proteolysis into the systems realm. Previously, the function and specificity of the catalytic cleft of many classes of proteases has been studied with: 1) synthetic peptide libraries [17, 18], 2) covalent active site probes and suicide substrates [19, 20], 3) substrate phage display [10, 21] and 4) proteome-derived peptide libraries [22]. Until now, no study has taken advantage of advances in new technology to gain the volumes of data that can instruct a systems view of proteolysis. Three recent studies have incorporated NGS into substrate phage profiling of the catalytic clefts of proteases [23-25], but approaches for the analysis of these large data sets to gain important mechanistic insight beyond what was possible with a typical substrate phage display experiment are lacking. Here, we describe a new strategy for interrogating protease function and are able to define the entire landscape of peptide substrate recognition, uncover for the first time unique sequences that are highly selective for closely related proteases, and estimate the contribution of the catalytic cleft to specificity of recognition of protein substrates.

## Results

### How many substrates define specificity of the catalytic cleft of a protease?

Enzyme specificity (defined as k_cat_/K_M_ for a given substrate relative to all others [26, 27]), is an elusive concept when applied to proteases, as more than one substrate can often have similar k_cat_/K_M_ for a given protease and more than one protease can have similar k_cat_/K_M_ values for the same substrates. As a result of this basic uncertainty, proteases are difficult to study [4]. Therefore, when describing specificity of proteases, it is useful to introduce a concept of “selectome”, which implies a multiplicity of substrates selective for a given protease. A selectome of a protease can be conceptually defined as a set of amino acid sequences of the length determined by the number and relative positions of selectivity determinants in its catalytic cleft, that only as a whole, is unique to that protease and thereby represents its proteolytic signature.

Protease specificity profiling is a combinatorial problem, i.e. the number of substrates in the selectome of a protease depends on the number of subsites as well as the number and stringency of selectivity determinants in its catalytic cleft. A protease with no selectivity determinants would cleave any peptide bond with equal efficiency and therefore has no definable selectome. A selective protease will have a pool of substrates with sequences distinct from those found in peptides with uniform distribution of residues across its length and thus reflective of the structure of its catalytic cleft. State of the art approaches available today allow only to probe the selectome of a given protease, but they are not designed to fully define it. More specifically, they allow to estimate the probability of observing a particular residue at a given position of a substrate based on the data sets obtained using peptide libraries [28, 29] or protein substrate databases (MEROPS). Unbiased selectivity profiling necessary to obtain the selectome of any protease relies on diversity of the sequence space used for that purpose. The closer to the theoretical maximum is the number of interrogated amino acid sequences in a library of potential substrates, the more accurate and detailed the resulting specificity profiles will be. In fact, it is implicit in all experiments aimed at unbiased profiling of substrate specificity that the results are representative of the catalytic cleft preferences. This means that it is assumed that the set of probes used for that purpose has a uniform distribution of amino acid patterns recognized by the catalytic cleft. This assumption, though basically important, is not routinely tested in experiments using synthetic and proteome-derived peptide libraries. Out of all existing approaches, only substrate phage display can meet the challenge of achieving maximum sequence diversity that can be quantitatively defined. The diversity afforded by substrate phage display, however, presents a technical challenge for determining the scissile bonds in millions of identified substrates. Combining Next Generation Sequencing (NGS) of substrate phage DNA with information theory-based data analysis allows to obtain highly defined selectomes without the need for experimental identification of scissile bonds. Due to the number of generated substrate sequences, this approach uniquely provides the possibility of quantifying specificity as well as selectivity and redundancy of proteases from the same phylogenetic families. In addition, detailed selectivity profiles obtained using this approach allow to determine how closely they represent the cleavage profiles derived from the protein substrates using N-terminomics and other experimental approaches. This, in turn, allows to establish if the catalytic cleft specificity is the main driver of physiologic substrate recognition or other features such as exosites or auxiliary domains are the primary determinants or modifiers of specificity.

When setting out to define the selectome of a protease, it is important to estimate how well its complexity is matched by the amino acid sequence library used for that purpose. While theoretically it may be possible to assess the weighted contribution of individual sites across the catalytic cleft to substrate specificity of a given protease with as few as 30 substrates of maximally variable composition [30], defining the selectome of that protease is a far larger combinatorial problem requiring a substrate library of appropriate diversity. Proteases vary widely in catalytic cleft specificity. An information entropy based analysis of specificity of proteases of all catalytic classes using the MEROPS database of protease substrates [11] demonstrates that while most serine, cysteine and aspartic proteases possess S1 (Schechter and Berger nomenclature, [31]) centric specificity, S3 and S1’ are the most stringent selectivity determinants in the catalytic cleft of metalloproteinases. The role of the S3 and S1’ sites in selectivity of Matrix Metalloproteinases (MMPs) is well documented [29, 32-34]. Since MMPs have two major selectivity determinants in their catalytic clefts that are located two subsites apart from each other, selectomes of these enzymes will range between 1 and 160,000 sequences spanning the P3-P1’ positions relative to the scissile bond (Fig. 1A). Sequence space covered by the randomized hexapeptide library, used in our phage display approach, (Fig. 1B, theoretical maximum = 6.4 × 10^7^) is adequate for interrogating selectome sizes of 160,000 and below. So, we used two closely related members (MMP-2 and MMP-9) of the 24-member Matrixin family and close to completely diverse library of hexapeptides displayed on gene 3 protein of M13 phage to explore and validate the concept of selectome.

**Figure 1.**
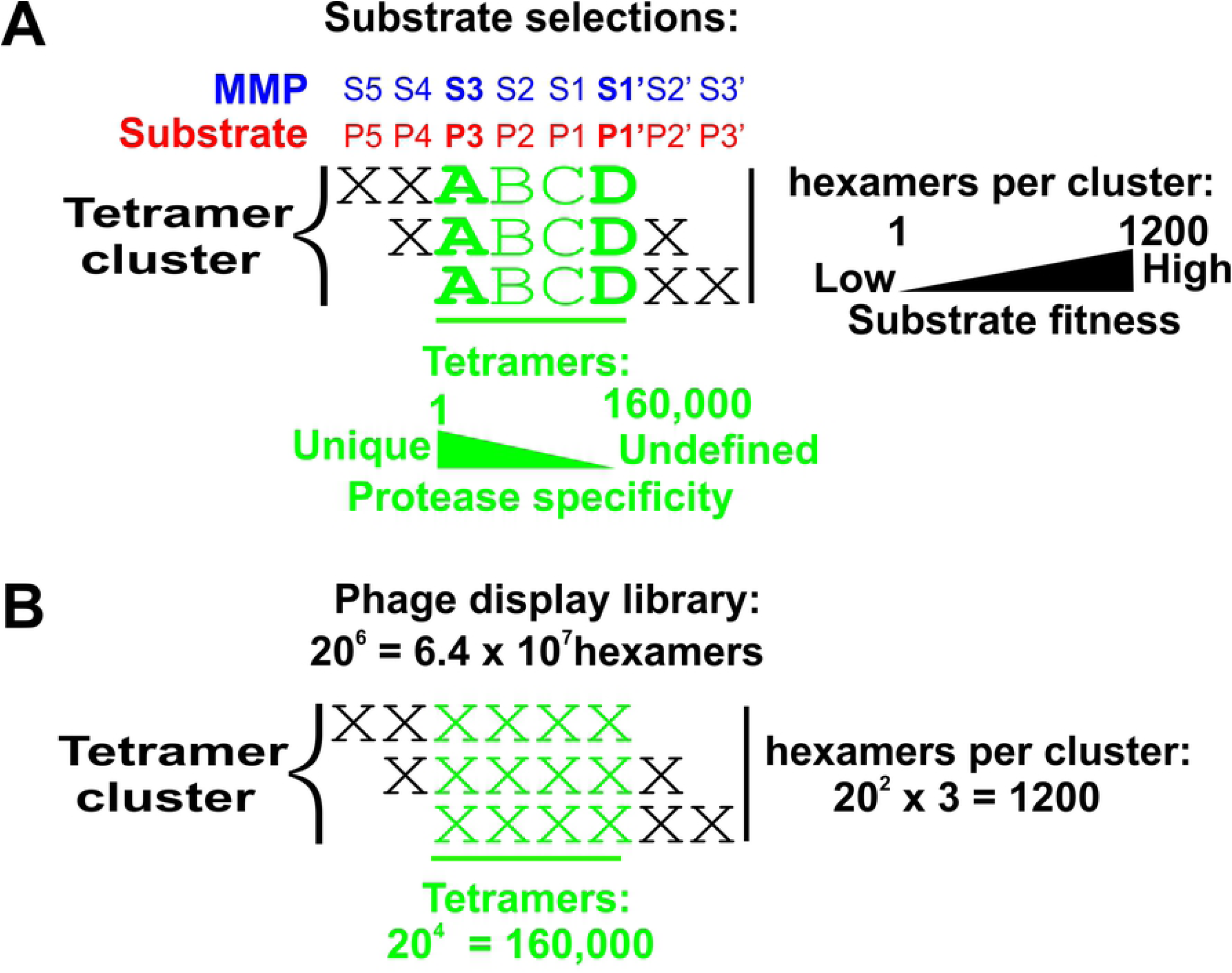
Basis for quantitative specificity profiling of the MMP catalytic cleft across S3 – S1’. **A**. *Interactions between individual subsites in the catalytic cleft of MMPs and the corresponding positions in the peptide substrates determine substrate fitness*. S3 and S1’ are the most selective binding sites in the catalytic cleft of the MMPs (bold red lettering). Together with the S2 and S1, they interact with P3-P1’ tetramers in substrates. Changes in repertoires of the P3 and P1’ residues in substrates (bold green lettering) affect substrate fitness the most. The fitter a particular tetramer is as a substrate, the larger the population of hexamers containing that tetramer in the substrate set (tetramer cluster) will be. The number of amino acid hexamers per tetramer cluster (1-1200) is a measure of fitness of a particular tetramer as a substrate. The larger the number of tetramer clusters in the substrate population (1-160,000) of a given protease, the less defined is the specificity of that enzyme. **B**. *Phage display library used for specificity profiling of MMPs has the diversity matching the theoretical maximum*. To interrogate specificity of MMP-2 and 9, we used a library of randomized hexapeptides displayed on gene 3 of M13 phage. The theoretical maximum of hexamer combinations is 6.4 × 10^7^. The theoretical maximum for the number of hexapeptides in a tetramer cluster is 1200. There are 160,000 combinations of natural amino acid residues in random tetramers.

The S3 and S1’ binding pockets along the catalytic cleft of MMPs together with S2 and S1 between them form a tetramer binding unit (Fig. 1A). The same tetramer combination of residues can occupy three different frames in a hexapeptide (Fig. 1B). When all the hexamers containing a given tetramer are present in the library, they will form a 1200-member tetramer cluster (Fig. 1B). The number of unique hexamer peptides containing the same tetramer (Fig. 1A) is a reflection of **substrate fitness** (related to k_cat_/K_M_) of a given tetramer sequence. If every hexamer peptide in a tetramer cluster can be found in the phage substrate set, then that tetramer has the substrate fitness and correspondingly the average k_cat_/K_M_ value at or above the threshold determined by the conditions of the experiment. Likewise, tetramer clusters with fewer than maximum numbers of hexamers per tetramer cluster must have lower than maximum substrate fitness and lower k_cat_/K_M_. The average k_cat_/K_M_ is defined as an average value of k_cat_/K_M_ over all hexamers belonging to the same tetramer cluster. The k_cat_/K_M_ threshold value is important to be able to compare fitness levels of substrates within the selectome of a given protease as well as across selectomes of different proteases. Choice of the k_cat_/K_M_ threshold value could be tricky, as ideally, it requires to have an estimate of the range of specificity constants for the protease being studied. We based our choice on the data previously published for the substrate phage display system used in this study [10, 33, 35]. The conditions for substrate phage selection were set so that 99% of all substrates with k_cat_/K_M_ of 3,289 M^-1^s^-1^ would be cleaved. This means that it would take 200 nM protease 2 hours to digest 99% of substrates with the k_cat_/K_M_ of 3,289 M^-1^s^-1^, provided the substrate concentration is much below K_M_, which is the case in our experiment. Substrates with k_cat_/K_M_ values above the threshold will be cleaved faster and those below slower.

We performed two rounds of selection of MMP-2 and 9 substrates using the conditions described above (see Materials and Methods for details). Sequences of hexamer peptide substrates were obtained using NGS analysis of the substrate phage DNA (See Supplemental data for details). Next, we grouped each of the hexamer peptides containing the same tetramer sequences into tetramer clusters (Fig. 1A, S1 Table, S3 Table, S4 Table). In order to avoid influences of hexamers belonging to more than one tetramer cluster and potentially representing co-occurring cleavages in the same hexamer or positions outside of the P3-P1’ tetramer, each hexamer MMP substrate was assigned to the most abundant tetramer cluster it can be found in. Thus, every substrate hexamer belongs to a single tetramer cluster. In parallel, we have also performed NGS analysis of the naïve (or initial) phage display library and also grouped the hexamer peptide sequences into tetramer clusters (Fig. 1B, S1 Table, S2 Table), but without eliminating redundancy (i.e. allowing the same hexamer to belong to more than one tetramer cluster), since no selective pressure that could influence the distribution of the tetramer clusters has been applied in generating this set. Next, we compared the distributions of relative abundances of tetramer clusters in the naïve phage display library and MMP-2 and 9 substrate selections. As can be seen in Fig. 2A, the distribution of relative abundances of tetramer clusters has changed significantly following substrate selections. MMP substrate sets have much broader distributions of relative abundances than the naïve library with the majority of tetramer clusters having lower probabilities than those in the naïve library. To determine the degree of change between the distributions of tetramer cluster probabilities in the substrate selections relative to the naïve library, we calculated the Shannon entropy values for each of the distributions:

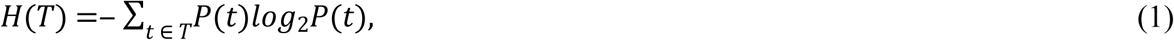

where *H(T)* is Shannon entropy of the distribution of tetramer clusters *(T)* each with probability *P(t)*. The probability of a given tetramer can be defined as:

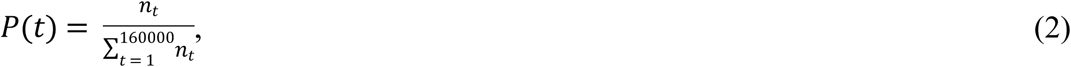

where *P(t)* is the probability of the *t*^*th*^ tetramer calculated as the ratio between the number of hexamers in that tetramer cluster *(n*_*t*_) and the number of hexamers in the entire set of tetramer clusters (160,000 tetramers). The naïve phage display library has the Shannon entropy value of 17.218 (S5 Table, S6 Table, which is not very far from that of a uniform distribution of tetramer clusters equal to 17.288 (log_2_ 160,000). This is an important characteristic of the library we used for substrate selections that gives an idea of its diversity relative to the maximum. Shannon entropy values of the substrate sets are 13.93 and 13.67 for MMP-2 and 9, respectively (S5 Table, S6 Table), which are significantly lower than that of the naïve library, as expected based on the changes in the probability distributions between the naïve library and the substrate selections (Fig. 2A).

**Figure 2.**
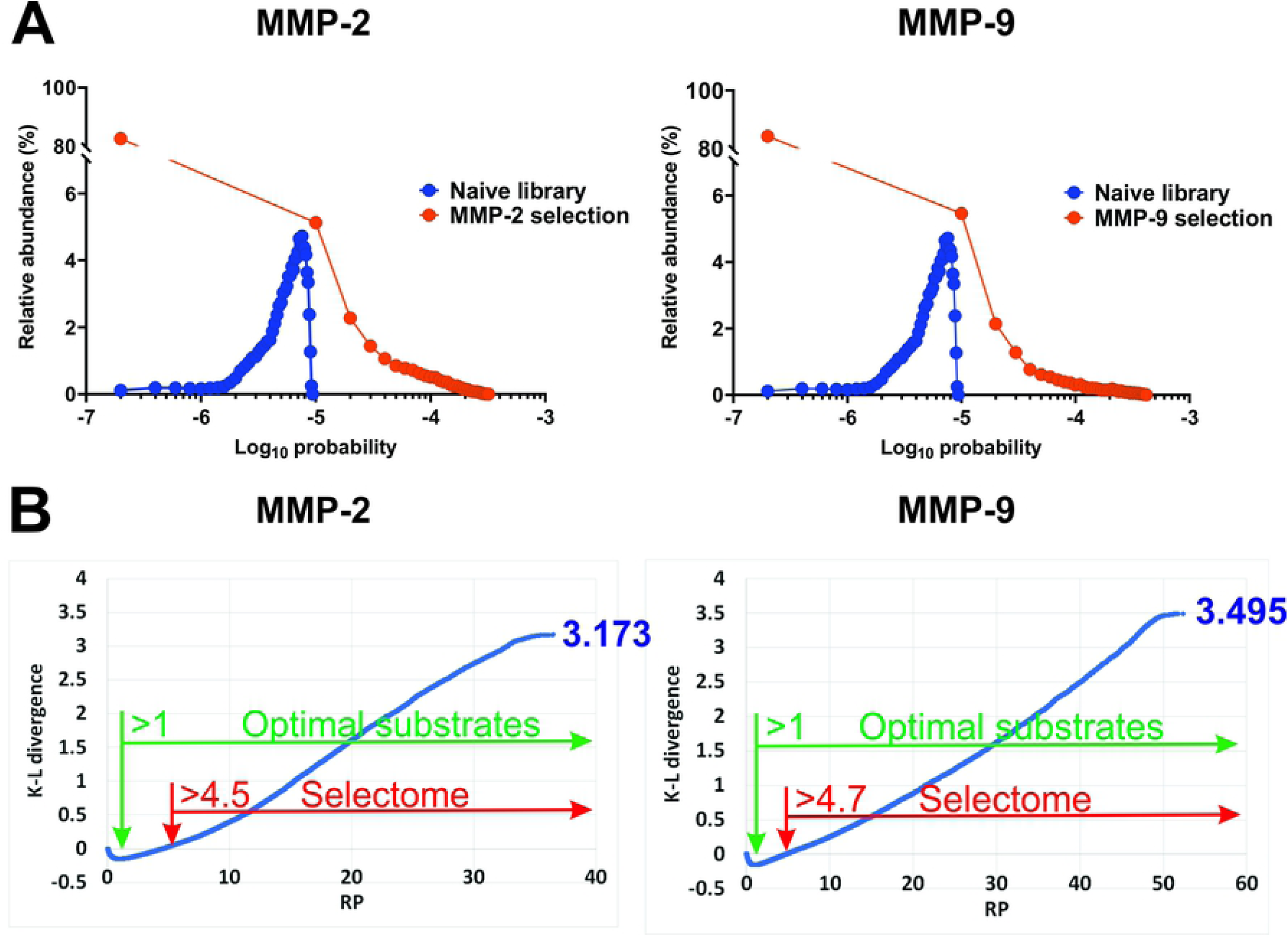
Divergence between probability distributions of tetramer clusters in substrate selections and the naïve library is a measure of substrate specificity. **A**. *Distributions of relative abundances of tetramer clusters in the MMP substrate selections are significantly different from that in the naïve phage display library*. Tetramer clusters within evenly spaced ranges of probabilities were binned together and their relative abundances were plotted as a function of log_10_ of average probabilities in the respective sets. **B**. *A subset of tetramer clusters in substrate selections with positive cumulative contribution to K-L divergence relative to the naïve library constitutes the selectome of a protease*. Cumulative contribution of individual tetramer clusters with RP values between of 0 and 4.5 for MMP-2 and 0 and 4.7 for MMP-9 to the K-L divergence relative to the naïve library is equal to 0. The K-L divergence between the probability distributions in the substrate sets of a protease with no definable specificity and the naïve library is always equal to 0. Therefore, the substrate sets with overall positive cumulative individual contributions to K-L divergences constitute the selectomes of proteases with definable specificities (MMP-2 and 9, red arrows). Tetramer clusters with RP values above 1 have positive individual contributions to the K-L divergences of the substrate selections and, therefore, are considered optimal substrates (green arrows).

To characterize the distribution of probabilities of tetramer clusters in the substrate sets in terms of substrate fitness related to the catalytic efficiency, we introduced the ratio between probabilities (Relative Probability or RP) of finding a given tetramer cluster in the MMP selections relative to the naïve phage display library, which can be used as a measure of substrate fitness of a tetramer relative to all others in a substrate set:

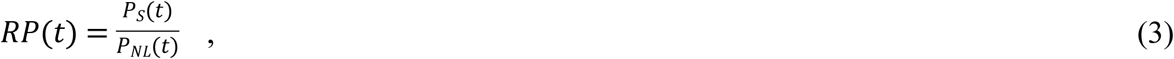

Where *P*_*S*_*(t)* is the probability of a *t*^*th*^ tetramer cluster in substrate selection and *P*_*NL*_*(t)* is the probability of that tetramer cluster in the naïve phage display library. Furthermore, the use of RP eliminates potential biases in tetramer probability distributions of the substrate sets due to deviation from uniformity of the tetramer probability distribution in the naïve library and potential differences in sequencing depths between the two sets.

The RP value for tetramer clusters in substrate sets has a theoretical range of maximum values between 1 (for a non-specific protease cleaving all tetramer substrates with equal efficiency: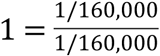) and 160,000 (for a maximally specific protease with only one tetramer substrate: 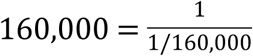). Importantly, RP must correlate with relative substrate fitness across its range of values for a given protease. To validate this assumption, we used the data obtained for a published set of 1369 phage substrates with experimentally determined scissile bonds and K_(obs)_ values (S7 Table) [10]. In this set, of all substrates containing non-redundant tetramers only 1.2% and 1.9% had no matching tetramer clusters in the MMP-2 and MMP-9 substrate selections, respectively. This observation confirms accuracy of the P3-P1’ assignments in tetramer clusters of the substrate selections. Next, we performed a standard statistical binary classification test (S8 Table) at increasing values of RP to determine if RP is a good predictor of a phage displayed hexamer peptide being a substrate. In this analysis, all P3-P1’ tetramers with RP values above a certain threshold and a non-zero value of K_(obs)_ were considered as true positives (TP). All tetramers with RP values below that threshold and a K_(obs)_ equal to 0 were considered as true negatives (TN). If a value of RP was above the threshold, but the K_(obs)_ was equal to 0, then the tetramer was classified as a false positive (FP). Finally, the tetramers with RP values below a threshold but a non-zero K_(obs)_ were classified as false negatives (FN). We expected the prediction to improve as the RP threshold value increased. Indeed, the Mathews Correlation Coefficient (MCC) improved significantly when the RP threshold value increased from 0 to 1, mostly due to a decrease in the FP and an increase in TN rates, respectively (S8 Table). Further increase in RP threshold value did not result in a significant change in MCC but the FP and the TN rates continued to decline. This analysis indicates that there is a positive correlation between K_(obs)_ of individual phage substrates and RP of the matching P3-P1’ tetramers.

Next, we performed an analysis of correlation between RP of tetramer clusters and K_(obs)_ of the hexamer substrates containing the matching P3-P1’ tetramers. It is expected that only averages of the catalytic efficiency constants of hexamer substrates contributing to a particular tetramer cluster will correlate with the RP value of that tetramer cluster because residues outside of the P3-P1’ tetramer will affect the catalytic efficiency to some degree. It is impractical to determine the K_(obs)_ value of each of the hexamer substrates in each of the tetramer clusters to obtain the averages. So, we used the following approach to make the correlation analysis feasible. First, we obtained the RP values for the tetramer clusters matching the P3-P1’ positions in hexamer substrates. Next, the tetramers were grouped based on their RP values to generate an evenly spaced distribution of bins across the RP range. The bins were distributed either 4 or 5 units apart across the RP range The average values, and standard errors of the RP and corresponding K_(obs)_ have been calculated for substrates in each bin. As shown in Fig. 3A, linear regression analysis demonstrates that average RP significantly correlates with average K_(obs)_ for phage substrates of both MMP-2 and 9.

**Fig. 3.**
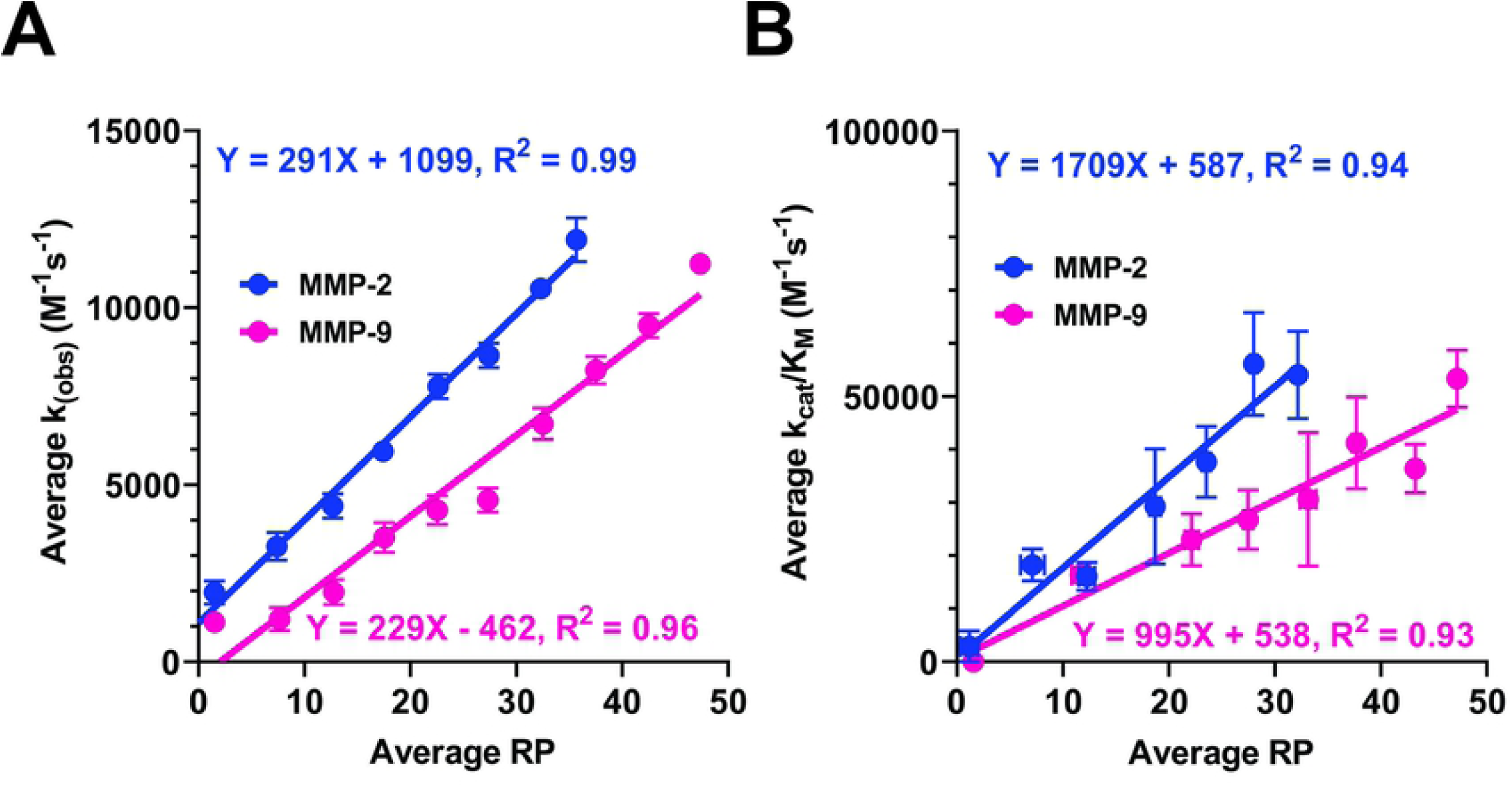
Probability of finding a tetramer cluster in substrate selections relative to the naïve library (RP) correlates with substrate fitness. K_(obs)_ values for 1369 individual phage substrates or k_cat_/K_M_ values of 100 peptides derived from substrate phage sequences were experimentally determined as described in the text. The substrates were binned into evenly distributed groups based on the RP values of the tetramer clusters corresponding to their P3-P1’ positions in substrates. The average K_(obs)_ (**A**) or k_cat_/K_M_ (**B**) values for each bin were plotted as a function of the corresponding average RP values and the data were subjected to linear regression analysis. The equations and goodness of fit parameters (R^2^) of the linear regression analyses of the MMP-2 and 9 data are shown at the top and bottom of the graph, respectively.

K_(obs)_ is a catalytic efficiency constant that was determined for each of the 1369 substrates in the set at 50 nM active enzyme for 2 hours and calculated using the integrated Michaelis–Menten equation [10]. This limits the range of the K_(obs)_ values that can be obtained to between 0 and 12,792 (M^-1^ s^-1^). So, all substrates with the true K_(obs)_ above this value will nevertheless have a K_(obs)_ equal to the preset maximum. With this limitation in mind, we corroborated the results using synthetic peptides, thereby extending the correlation to the entire range of k_cat_/K_M_ values for each MMP. Sequences of the 100 peptide set used for the analysis were derived from phage substrate selections and their k_cat_/K_M_ values were experimentally determined as described in [33] (S9 Table). The correlation analysis was carried out the same way as for the phage substrates (Fig. 3B). The results clearly demonstrate that the average k_cat_/K_M_ values of hexamer substrates correlate with the average RP values of the tetramer clusters they belong to. This validates the use of RP as a measure of substrate fitness.

Substrate selections contain close to half of theoretically possible tetramer clusters, most of which have RP values below 1. Distributions of the RP values in substrate selections of MMP-2 and 9 do not reach a plateau at the high end (S2, Table, S3, Table). Therefore, these probability distributions must be representative of the entire ranges of the respective catalytic cleft specificities. Since the majority of the tetramer clusters in the substrate sets of MMP-2 and 9 have probabilities lower than in the naïve library and constitute rare events, they must be relatively poor substrates and therefore contribute little if at all to the specificity of the two enzymes. We asked a question if one could find an appropriate threshold to select the tetramer substrates with statistically significant contribution to specificity of the catalytic cleft. To that end, we used Kullback-Leibler (K-L) divergence [36] as a measure of distinction between the probability distributions of tetramer clusters in substrate selections and the naïve phage display library, thus reflecting specificity of the catalytic cleft:

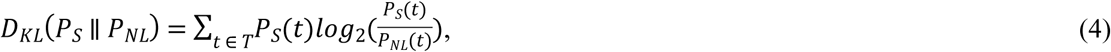

Where *D*_*KL*_ *(P*_*S*_‖*P*_*NL*_) is the K-L divergence between the tetramer probability distributions in the selections *P*_*S*_*(t)* and the naïve library *P*_*NL*_*(t)* defined on the same probability space *T*.

The K-L divergence, or relative entropy determines how one probability distribution is different from another, reference distribution. The larger the value of relative entropy, the more divergent the probability distributions of the test and reference sets are. The K-L divergence values can range from 0 for a protease with no specificity to 17.288 for a perfectly specific protease with a single tetramer substrate assuming uniform probability distribution for the reference set. We performed K-L divergence analysis using probability distributions of tetramer clusters in the MMP selections as the test and those in the naïve library as the reference sets, respectively. The relative entropies are 3.173 and 3.495 (S5 Table and S6 Table) for MMP-2 and 9 tetramer clusters, respectively, indicating that MMP-9 has a narrower specificity than MMP-2, although not by much. The total number of tetramer clusters with non-zero probabilities *P(t)* and thus non-zero contributions to the values of K-L divergence, is 78,757 and 76,696 for MMP-2 and 9, respectively. The plots of the sum of individual components in the calculations of the expected value using equation 4, as a function of RP for MMP-2 and 9 have two distinct parts: one below and the other above the zero value of K-L divergence (Fig. 2B). While the former has no net contribution to K-L divergence, the latter is the sole contributor. The RP value at the intersection of the line in the graph with the X-axis is a useful threshold to define the set of tetramer clusters, which as a whole, is unique to a given protease and therefore represents its “**selectome**”. These values are 4.5 and 4.7 for MMP-2 and 9, respectively (indicated by red arrow in Fig. 2B). There are 7,921 and 6,094 tetramers above the RP threshold, belonging to the MMP-2 and 9 selectomes, respectively (S5, S6 Tables). They constitute 8-10% of all tetramers with non-zero value of RP. Another useful threshold, at which the probabilities of finding a tetramer in the substrate selection and in the naïve library are the same, occurs at the RP value of 1. Tetramer clusters with the RP values greater than 1 are considered optimal substrates (marked by the green downward arrow in the Fig. 2B), since their individual contributions to the K-L divergence are positive. The numbers of tetramers corresponding to the RP values above 1 are 16,395 in the MMP-2 and 15,581 in the MMP-9 substrate sets. Tetramer clusters with RP values less than 1 are considered suboptimal substrates, since they contribute negatively to the K-L divergence. Thus, out of the total of 78,757 MMP-2 and 76,696 MMP-9 non-zero tetramer clusters (i.e. containing at least one hexamer) found in the selection sets, ∼20% are optimal substrates. Based on the statistically defined threshold introduced above, the selectome constitutes a unique subset of substrates of a given protease.

To corroborate the findings of the K-L divergence analysis, we looked at the distributions of tetramer clusters across the RP range in 10% increments from highest to lowest (S1(A) Fig.). The number of tetramer clusters across the RP range shows a slow increase until it reaches the lowest 10%, when it increases dramatically. S1(B) Fig. shows the distribution of the numbers of hexamers per tetramer cluster across the same intervals. Not surprisingly, the lowest 10% have a precipitous decline in that metric compared to the nearest neighbor. Thus, this analysis of tetramer cluster distributions agrees with the relative entropy-based analysis, showing that the top 10% of tetramer clusters are populated the highest, which is consistent with percentages of tetramer clusters in the selectomes of MMP-2 and 9. To put these data in perspective, one must keep in mind that the tetramer clusters with positive cumulative contribution to K-L divergence (the selectome) in the set of MMP-2 substrates contain 2.31 × 10^6^ hexamers substrates, while those with zero cumulative contribution to K-L divergence, (RP interval between 0 and 4.5) – only 0.56 × 10^6^. The same numbers for MMP-9 are 1.64 × 10^6^ and 0.54 × 10^6^, respectively. So, 80% of hexamer substrates of MMP-2 and 75% of MMP-9 belong to their respective selectomes. This observation provides basis for the conclusion that the catalytic cleft specificity of MMP-2 and 9 is primarily defined by S3-S1’ subsites, as expected. The poorly populated tetramer clusters are represented by sequences that, as P3-P1’ tetramers, are suboptimal substrates, whose fitness may be modulated by exosites outside S3-S1’ and which are found in the minority (20-25%) of the hexamer substrates.

In this section of the results we have developed a concept of substrate specificity we call “selectome”, which though intuitive, is not easy to grasp. To the best of our knowledge, there have been no prior reports of an approach aimed at defining the set of substrates that fully captures substrate specificity of a protease. In the following sections, we will substantiate this concept by applying it to analyses of catalytic cleft selectivity between MMP-2 and 9 and contribution of the catalytic cleft specificity to protein substrate recognition.

### Quantifying substrate specificity

Distribution of tetramer clusters across the normalized RP (RP/RP_Max_) spectrum (Fig. 4A) can be used to quantify the catalytic cleft specificity in terms of both substrate numbers and fitness. A protease with no specificity will have all possible 160,000 tetramer clusters with RP/RP_Max_ value of 1, since it can cleave all peptide bonds regardless of context with equal efficiency. A perfectly specific protease will have just one tetramer cluster with RP/RP_Max_ value of 1, since this is the only tetramer it can cleave. Specificity can be approximated by the **Total Substrate Fitness** across the RP range (TSF). TSF is numerically equivalent to the area under the curve representing the number of tetramer clusters as a function RP/RP_Max_. We used a simple trapezoidal formula for numerical integration to calculate TSF.

**Figure 4.**
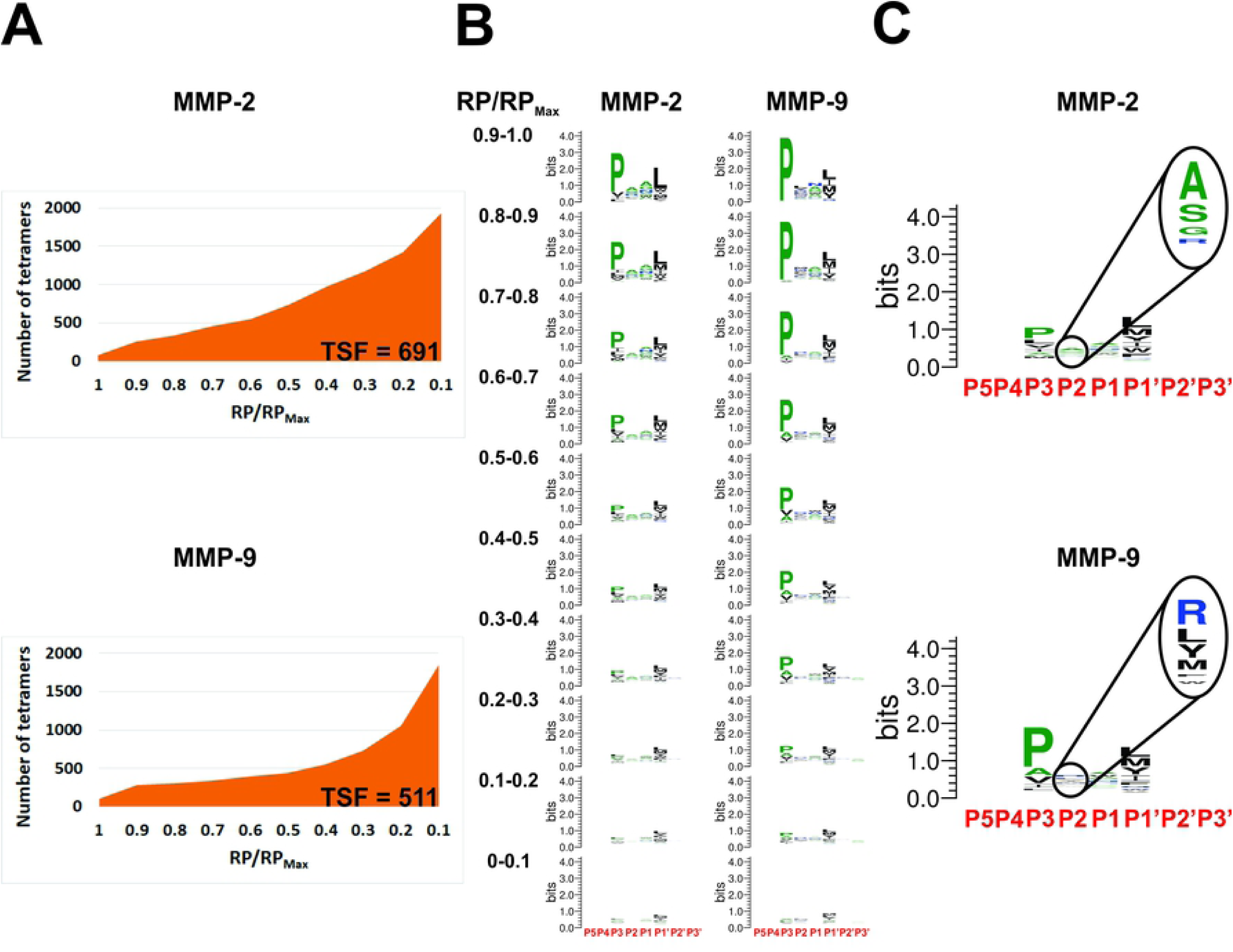
Total substrate fitness and composition of the selectomes of MMP-2 and 9 reflect differences in their specificities. **A**. *Substrate number and fitness can be used for quantification of substrate specificity*. Total Substrate Fitness (TSF) of a protease is calculated as area under the curve of the plots of substrate number as a function of relative substrate fitness (RP/RP_Max_). **B**. *Selectome-based specificity profiles of MMP-2 and 9 reflect changes in substrate composition as a function of substrate fitness*. Peptide hexamers in the selectomes of MMP-2 and 9 were aligned along the P5-P3’ positions based on P3-P1’ matches in the corresponding tetramer clusters and divided into 10 groups according to their RP values relative to the maximum (RP/RP_Max_). Relative abundances of amino acid residues at each position were calculated and presented in the form of a logo plot for each of the groups. **C**. *Aggregate specificity profiles of MMP-2 and 9 reveal the major selectivity features of MMP-2 and 9*. Hexamers belonging to the tetramer clusters constituting the selectomes of MMP-2 and 9 were aligned along the P5-P3’ positions based on P3-P1’ matches in the corresponding tetramer clusters, and relative abundances of residues at each position were plotted in the form of a logo. Zoom-in ovals show the relative contributions of residues to the P2 specificity profile of each enzyme.

A non-specific protease will have 1.6 × 10^5^ tetramer clusters times the RP/RP_Max_ value of 1, which yields a TSF value of 1.6 × 10^5^ (Fig. 1B). A perfectly specific protease will have 1 tetramer cluster times the RP/RP_Max_ value of 1 yielding a TSF value of 1. According to this calculation, TSF of MMP-2 and 9 selectomes is equal to 691 and 511, respectively (Fig. 4A). Thus, the catalytic cleft specificity of MMP-9 is narrower and constitutes 74% of that of MMP-2.

To analyze the composition of the MMP-2 and 9 substrates in respective selectomes, hexamers comprising the P3-P1’ tetramer clusters were aligned along the P5-P3’ interval and sequence logos were generated based on frequency of occurrence of residues at individual positions using a sequence logo generator WebLogo [37]. Compositions of the tetramer clusters in the selectomes of MMP-2 and 9 are shown in Fig. 4B and C. Consistent with the roles of S3 and S1’ as primary selectivity determinants in the catalytic clefts of MMPs, P3 and P1’ positions in substrates contribute the most to substrate specificities of both MMP-2 and 9. Logo plots of distributions of residues along the P5 -P3’ (Fig. 4B) interval show dominance of the P3 position in tetrameric clusters with highest relative probabilities, which diminishes together with RP. Relative contribution of the P1’ position to the information content of the logo plots grows as relative probability decreases. These findings suggest that in general, the P1’ position of MMP substrates determines whether a particular sequence will be cleaved and the P3 position primarily determines the substrate fitness of a given tetramer sequence provided the P1’ content is compatible with the S1’ pocket specificity. Patterns of amino acid distribution across P3-P1’ positions of MMP-2 and 9 selectomes (Fig. 4C) are similar in general to those published elsewhere for substrates of MMP-2 and 9 [10, 29, 38]. However, it is striking to see that two datasets in separate publications from the same group have rather different compositions of residues in the P1’ position of substrates obtained from proteome-derived tryptic peptide libraries. The P1’ position of substrates in the dataset published in [22] has the variety of residues very similar to that observed in the MMP-2 selectome obtained by us. In the dataset published in [29] the P1’ position of the substrates is overwhelmingly occupied by Leu and Ile, with unnoticeable contributions from Met, Trp, Phe and Val, which are clearly present in the former dataset and also in the MMP-2 selectome obtained by us. This is evidence of experimental variability or discrepancies in diversity between the peptide libraries used in each case. Our data in Fig. 4B show a clear distinction between the aggregate specificity profiles of MMP-2 and 9 at the P2 position of substrates. The P2 repertoire of MMP-9 is very different from MMP-2, with significant contributions of residues with aliphatic (Leu and Met) and aromatic (Phe, Tyr, Trp) side chains compared to MMP-2’s Ala, Ser and Gly. There is structural basis for S2 selectivity between MMP-2 and 9 which has already been reported in [39] and will be discussed further in the text. The MMP-9 dataset published in [29] is strikingly similar to that of MMP-2 in the same study with P2 repertoire consisting of Ala, Ser, Met and Ala, Ser, Gly, respectively, which contradicts other findings. Not surprising is the absence of P2 Arg in the MMP2 and 9 substrates of their datasets and its presence in ours, since trypsin was used to generate the proteome derived peptide library. Interestingly, while the profiles of MMP-2 and 9 at the selective P2 position are different between our data set and that published in [29], the P1 profiles invariant between MMP-2 and 9 are very similar in both data sets. This suggests that the peptide library used in [29] was underrepresented in sequences selective for MMP-9 over MMP-2.

In this section we have introduced for the first time, a measure of protease specificity that reflects both substrate numbers and their relative catalytic efficiencies. We have also shown how substrate composition changes across the fitness range, which provides valuable insight into the correlation between catalytic efficiency and subsite specificity.

### Comparative analysis of selectomes reveals distinctions between selectivity determinants of MMP-2 and 9

One of the central obstacles to understanding protease biology is functional redundancy and specificity overlap between proteases from the same phylogenetic groups [4]. Selectome profiling presented in this study makes it possible to determine how much overlap and distinction there is between specificities of closely related proteases. Catalytic domains of human MMP-2 and 9 are 73% identical and 81% similar in their amino acid sequences. Direct comparison reveals that out of the total of 10,110 tetramers comprising the combined selectomes, 3,902 are shared by both, and 4,019 and 2,189 are found exclusively in the respective selectomes of MMP-2 and 9 (Fig. 5A, S10-S13 Tables). Thus, a pair of 73% identical proteases has only 39% of the combined selectomes in common, demonstrating a significant amount of S3-S1’ distinction between the two MMPs. MMP-2 has the broader specificity of the pair with 40% unique tetramers, while MMP-9 has only 22%, almost two-fold less than its closest relative in the MMP family.

**Figure 5.**
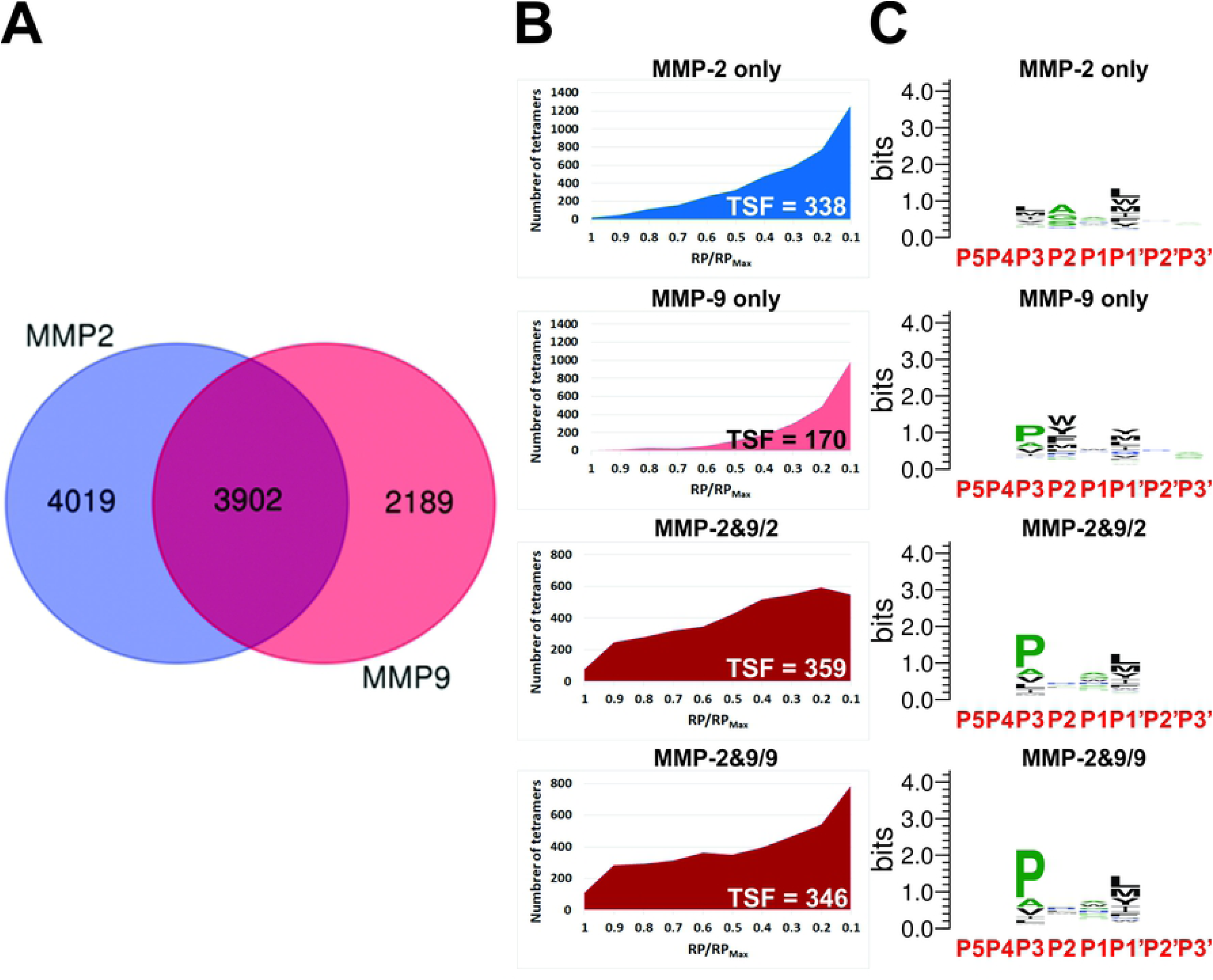
Comparative analysis of selectomes reveals divergent and conserved features in substrate recognition between MMP-2 and 9. **A**. *Venn diagram of substrate specificity overlap and distinction between the selectomes of MMP-2 and 9 shows significant selectivity between the two enzymes*. Tetramer clusters comprising the selectomes of MMP-2 and 9 were grouped based on their occurrence in the individual and overlapping substrate sets and the corresponding numbers are shown in the Venn diagram. For making Venn diagram we used on-line service at: http://bioinformatics.psb.ugent.be/webtools/Venn/. **B**. *Total Substrate Fitness (TSF)-based quantification shows that MMP-9 is more selective than MMP-2*. Numbers of tetramer clusters were plotted as a function of relative substrate fitness (RP/RP_Max_) to determine the TSF values of the unique and overlapping substrate sets of MMP-2 and 9 selectomes. Distributions of tetramer clusters across the relative RP range are different in the overlapping set of the two enzymes, hence denoted as MMP-2&9/2 for MMP-2 and MMP2&9/9 for MMP-9. **C**. *Aggregate specificity profiles based on the unique and overlapping tetramer clusters of MMP-2 and 9 reveal the distribution of selectivity across the catalytic cleft*. Hexamer peptides belonging to the tetramer clusters constituting the selective and common substrate sets of MMP-2 and 9 were aligned along P5-P3’ positions based on the P3-P1’ matches in respective tetramers and the relative abundances of residues at each position were plotted as logos.

Substrate fitness distributions of the selective and common sets of tetramer clusters across the RP range is shown in Fig. 5B. Total substrate fitness calculations show that selectivities of MMP-2 and 9 are equal to 338 and 170, respectively. Based on this, MMP-2 is 2-fold less selective than MMP-9, which is consistent with the numbers of unique tetramer clusters between the two selectomes. TSF values of substrates found in overlapping selectomes of MMP-2 and 9 are very similar, as expected, and equal to 359 and 346, respectively.

Logo profiles based on the selective tetramer clusters (S2 Fig.) show consistent patterns across the RP range with decreasing information content as the RP value goes down, as expected. The logos representing all substrates from the selective tetramer clusters (Fig. 5C) show the aggregate picture across the entire range of RP. Both MMP-2 and 9 display selectivity at S3, S2 and S1’ subsites of the catalytic cleft with similar contributions from each. Very different, however, are the repertoires and relative abundances of residues in substrates at the positions reflecting selectivity of each of the subsites. Previously, we have mapped the selectivity determining positions (SDPs) in the catalytic cleft of the MMP family [10]. By comparing the compositions of the subsites contributing to selectivity between MMP-2 and 9, one can account for distinctions observed between the unique substrate sets (Fig. 6). SDPs at S4/S3 junction (Gly175 in MMP-2 and Gln199 in MMP-9) and S2/S3 junction (Ala179 in MMP-2 and Pro192 in MMP-9) control the height of the catalytic groove at the S3 position together with the conserved S3 Tyr155/179 in MMP-2 and MMP-9, respectively. Distances controlling the height of the catalytic cleft in MMP-2 are 11.4 Å between Tyr155 and Gly175 and 11.1 Å between Tyr155 and Ala179. In MMP-9 the space between the corresponding residues is narrower (9.3 Å between Tyr179 and Gln199 and 9.6 Å between Tyr179 and Pro192), which is consistent with the presence of the large aliphatic Leu, Met and Ile in the P3 positions of the MMP-2 selective substrates and their virtual absence in the MMP-9 ones. More compact Pro and to a lesser extent Ala and Val at the P3 of the substrates are accepted by MMP-9.

**Figure 6.**
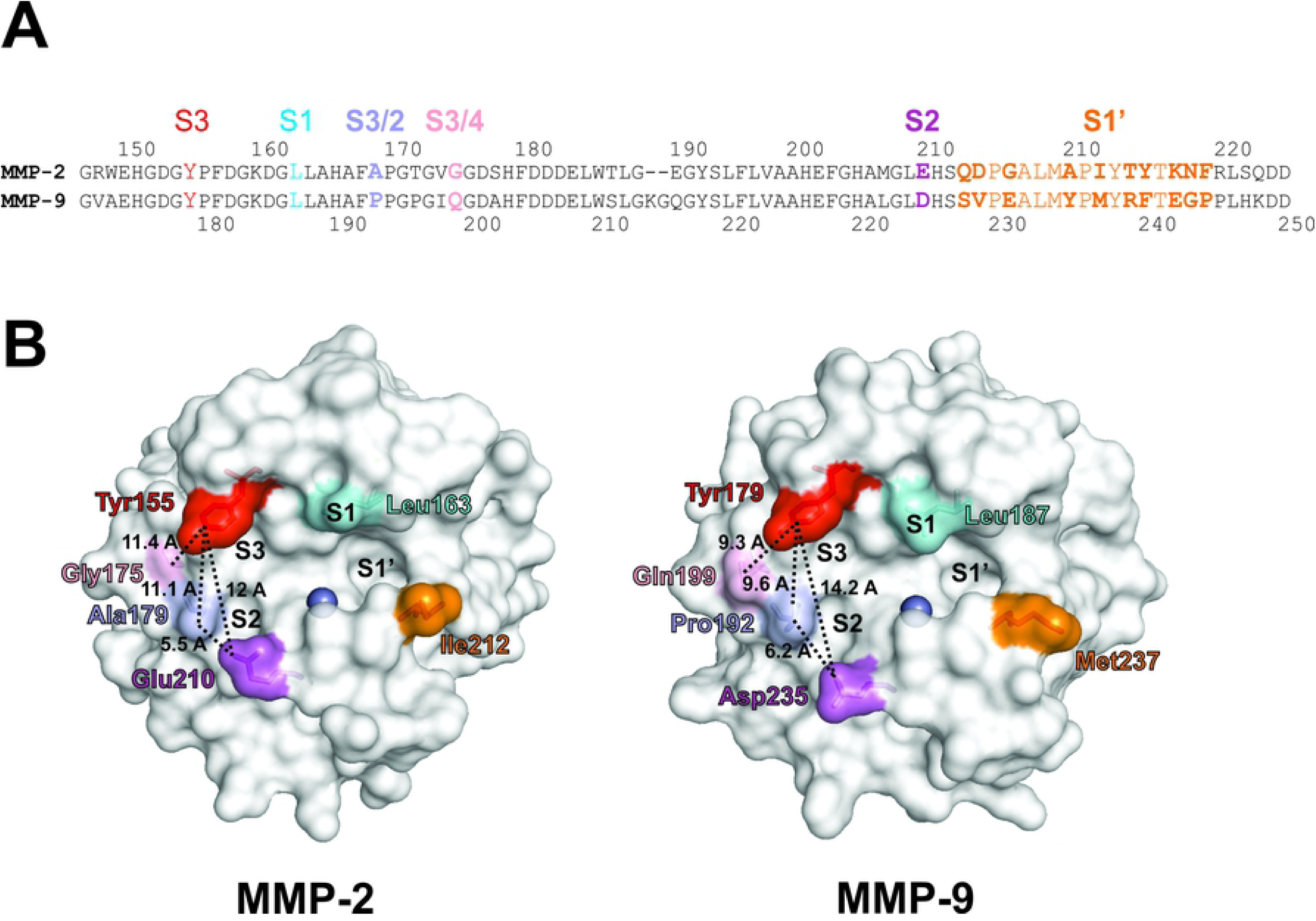
Changes in composition of selective substrates correlate with changes in determinants of selectivity between MMP-2 and 9. **A**. *Distribution of the selectivity determinants across the subsites of the catalytic clefts of MMP-2 and 9*. Sequences of the catalytic domains of MMP-2 and 9 were aligned based on the crystal structures of the catalytic domains of the respective enzymes (PDB IDs: 1QID for MMP2, 1GKC for MMP9). Residue numberings from the native N-termini are shown above (MMP-2) or below (MMP-9) the sequence. Selectivity Determining Positions (SDPs) are shown in color and marked in larger font. The catalytic cleft subsites they contribute to are shown directly above each SDP. Residues marked in bold differ between the two proteases (see text for details). **B**. *Structural features of the SDPs in the catalytic clefts of MMP-2 and 9 provide basis for experimentally determined subsite selectivity*. Residues contributing to the SDPs at S3, S2, S1 and S1’ binding pockets are shown on the surface representations of the three-dimensional structures of MMP-2 and 9 in colors matching the sequence alignments in A. See text for more details PyMOL molecular visualization system was used for display and analysis of 3D structures.

Another notable difference between the MMP-2 and 9 specificities is evident from the repertoire of residues at the P2 position of the selective substrates, as discussed above. Dominance of Ala, Gly and Ser at the P2 of MMP-2 is contrasted by the preponderance of bulky aromatic and to some extent aliphatic side chain residues of the MMP-9 selective substrates (Fig. 4C, S2 Fig.). This observation is consistent with the differences in composition of the S2 binding pocket sandwiched between the S2/S3 Ala179 and S2 Glu 210 in MMP-2 and S2/S3 Pro192 and S2 Asp 210 in MMP-9 (Fig. 6). The distance difference between these residues is 5.5 Å in MMP-2 A vs. 6.2 Å in MMP-9, in agreement with the observed differences in P2 composition of the respective unique substrates. Additionally, bulkier Glu210 narrows the catalytic cleft in MMP-2 to 12 Å from 14.2 Å in MMP-9, which has a more compact Asp235 in that position (Fig. 6).

Quite remarkable is the lack of significant contribution of P1 to selectivity as expected based on identical residues at the S1 SDPs of both enzymes (Leu163 in MMP-2 and Leu187 in MMP-9).

Differences in P1’ composition of the selective tetramers are more difficult to explain structurally due to the complexity of the S1’ binding site, formed by an allosteric hydrophobic tunnel preferentially occupied by Leu, Trp, Met and Ile residues in the selective substrates of MMP-2. In MMP-9 selective substrates P1’ Leu, the preferred residue by the S1’ pocket of the entire MMP family, is virtually absent and becomes noticeable only in the lower (0.2-0.5) RP/RP_Max_ range of the MMP-9 tetramer clusters. Out of the 18 residues comprising the S1’ loop, 10 are different between MMP-2 and 9, with 5 non-conserved substitutions. The fact that the selective substrates of both enzymes have significant differences in the repertoires of the P1’ residues is consistent with significant differences in SDP compositions of the S1’ binding pocket between the two enzymes. Of note, however, is that one of the SDPs forming the opening of the S1’ tunnel is different between the two enzymes (Ile212 and Met237) and could be a significant contributor to selectivity.

Importantly, the composition of substrates comprising the overlapping set of 4,019 tetrameric clusters is consistent with the classic PxxL pattern common for most MMPs [10] (Fig. 5C). Interestingly, even though the combined specificity profiles of the tetrameric clusters shared by both enzymes are nearly identical, there are noticeable differences in the profiles of tetrameric clusters at different RP levels (S2 Fig.).

In this section, for the first time, we provided a quantitative measure of overlap and distinction between specificities of closely related proteases that accounts for both substrate number and fitness. In addition, comparative analysis of composition of the selectomes of MMP-2 and 9 presented here combined with information on location and composition of selectivity determinants provides clear structural basis for identifying SDPs responsible for selectivity between these closely related enzymes.

### Telling a target from a bystander: contribution of the catalytic cleft specificity to protein substrate recognition

One of the questions central to understanding protease function is how to distinguish between targeted and coincidental proteolytic events. It stands to reason that proteases and their physiologic substrates co-evolved to be integral parts of complex physiological processes [4, 40, 41]. Mechanisms underlying protease-substrate recognition involve exosite, auxiliary binding domain and catalytic cleft interactions. While the K_M_ value of a proteolytic event can be affected by interactions outside the catalytic cleft, the k_cat_ value is completely dependent on the substrate fitness around the scissile bond, which is related to the rate of formation of the transition state intermediate. If that rate is close to zero, a tight interaction outside the catalytic cleft will result in inhibition of the protease. High rate of formation of the transition state intermediate will result in faster hydrolysis if the K_M_ value is sufficiently high to allow for dissociation of the enzyme-product complex before the reverse reaction re-forms the enzyme-substrate complex. So, mechanistically, there is a fine balance that needs to be struck between the k_cat_ and K_M_ values for a physiologically relevant proteolytic event to be integrated into the larger context of underlying biology.

To assess the contribution of the catalytic cleft specificity to physiologic substrate recognition by MMP-2 and 9, we used the data on protein substrate hydrolysis obtained by us and those available in the literature. The data set published in [42] was taken as a benchmark for protein cleavage site identification due to the rigor of data analysis and independent verification (S14 Table). Based on the comparison of this data set with ours, 81 and 84% of all the cleavages in the MMP-2 and 9 substrate sets are optimal substrates. Of them, 71% of MMP-2 and 79% of MMP-9 cut sites belong to their respective selectomes (S14 Table). These numbers are not very far from the probability (86%) of unambiguous identification of cut sites of a protease with known specificity (Glu C) used by the authors for validation of the statistical model for cleavage site identification used in their study [42]. The rest of the identified cleavages (19% for MMP-2 and 16% for MMP-9) are either suboptimal substrates (RP<1, 13.6% for MMP-2 and 10.5% for MMP-9) or not substrates at all (RP=0, 5.5% for MMP-2 and 5.3% for MMP-9) as defined by our analysis. Thus, there is a very good correlation between a cut site being a part of the selectome of MMP-2 or 9 and also being a validated substrate of the same MMP.

In the publication we used as the benchmark [42], the criteria for cleavage site identification were set very stringently, so that the ratios between the iTRAQ reporter ion intensities in the MMP-treated samples and the untreated controls had to be ≥ 10 in order for N-terminally labeled peptides to meet the statistical threshold to be considered a candidate cleavage sites. This was done to achieve a reasonable compromise between the false positive and false negative rates based on the statistical model developed by the authors. To find out if the selectome-based classification can be applied to distinguishing between substrate and non-substrate N-termini, we performed N-terminomic analysis of MMP-2 and 9-treated secretomes derived from HEK293 cells [43]. Cells were cultured in serum-free medium in the presence of 1 µM GM6001 (a broad MMP inhibitor) to minimize background hydrolysis by endogenous MMPs. The secretomes were concentrated, buffer exchanged to remove the inhibitor and incubated with MMP-2 or 9 (750 nM for 2 hours) or left untreated and then labeled with TMT reagents (see Materials and Methods for details). Following denaturation and digestion with trypsin, the secretomes were subjected to offline HPLC fractionation and then analyzed by LC-MS/MS to reveal novel N-termini resulting from MMP treatment. S15 Table shows the entire list of N-terminally labeled peptides and the corresponding positions in annotated proteins arranged according to their log_2_ of the ratios of the TMT reporter ion spectral counts relative to the untreated controls (isotopic enrichment or IE). Overall, we identified 453 N-termini in 243 proteins in the MMP-2 and 1034 N-termini in 428 proteins in the MMP-9 treated samples. Plotting relative abundances of the N-terminal peptides with different RP values as a function of IE, expressed as multiples of standard deviation away from the average values for the entire sets, shows that the N-termini with RP values above the selectome thresholds for MMP-2 (RP>4.5) and MMP-9 (RP>4.7) are predominantly found in the intervals with σ > 1 above the average EI (Fig. 7). The further do the IE values go below σ = 1 the higher is the proportion of the N-termini with RP values below 1. Based on these observations, in our study, the IE cutoff for calling a labeled N-terminus a cleavage site resides one standard deviation above the population mean. To confirm that RP-based categorization is a good predictor of isotopic enrichment, we performed binary classification analysis of the data shown in Fig. 7. The results demonstrate that an RP value above the selectome threshold (RP=4.5 for MMP-2 and 4.7 for MMP-9) is the best predictor (MCC = 0.502 for MMP-2 and 0.435, respectively) of an N-terminal peptide to have an EI value above σ = 1 relative to the population mean (S16, Table). These data are highly consistent with what we observed using the benchmark data set discussed above and provide basis for distinguishing between the true positive and potential false positive cleavage site identifications. Proteomic analysis of protease substrates based on isotopic labeling of novel N-termini requires quantification of reporter ion intensities of the mass tag pair used for the control and protease-treated samples. This quantification only reports on the relative abundances of labeled peptides in each sample and therefore is subject to limitations. These include: 1) inaccurate reporting of peptides with more than one cleavage site, 2) wrong cut site identification due to C-terminal cleavage by protease of interest and a novel N-terminus resulting from an unrelated proteolytic event, 3) wrong cut site identification due to the probabilistic nature of the assignment of the N-terminal label if side chain labeling is present in the same peptide and 4) artefactually high isotopic enrichment for non-substrate labeled peptides due to experimental error or other unknown factors. The ability to classify the N-terminal peptides based on the RP value simplifies identification of cleavage sites regardless of isotopic enrichment, which makes candidate substrate identification much easier. Also, it allows to set aside potentially false identifications for further evaluation when a cut site has an RP value of 0. More importantly, knowing the RPs of tetramer clusters matching the cleavage sites is of considerable value for deciding if a particular cleavage event is a target or a bystander for a given protease.

**Figure 7.**
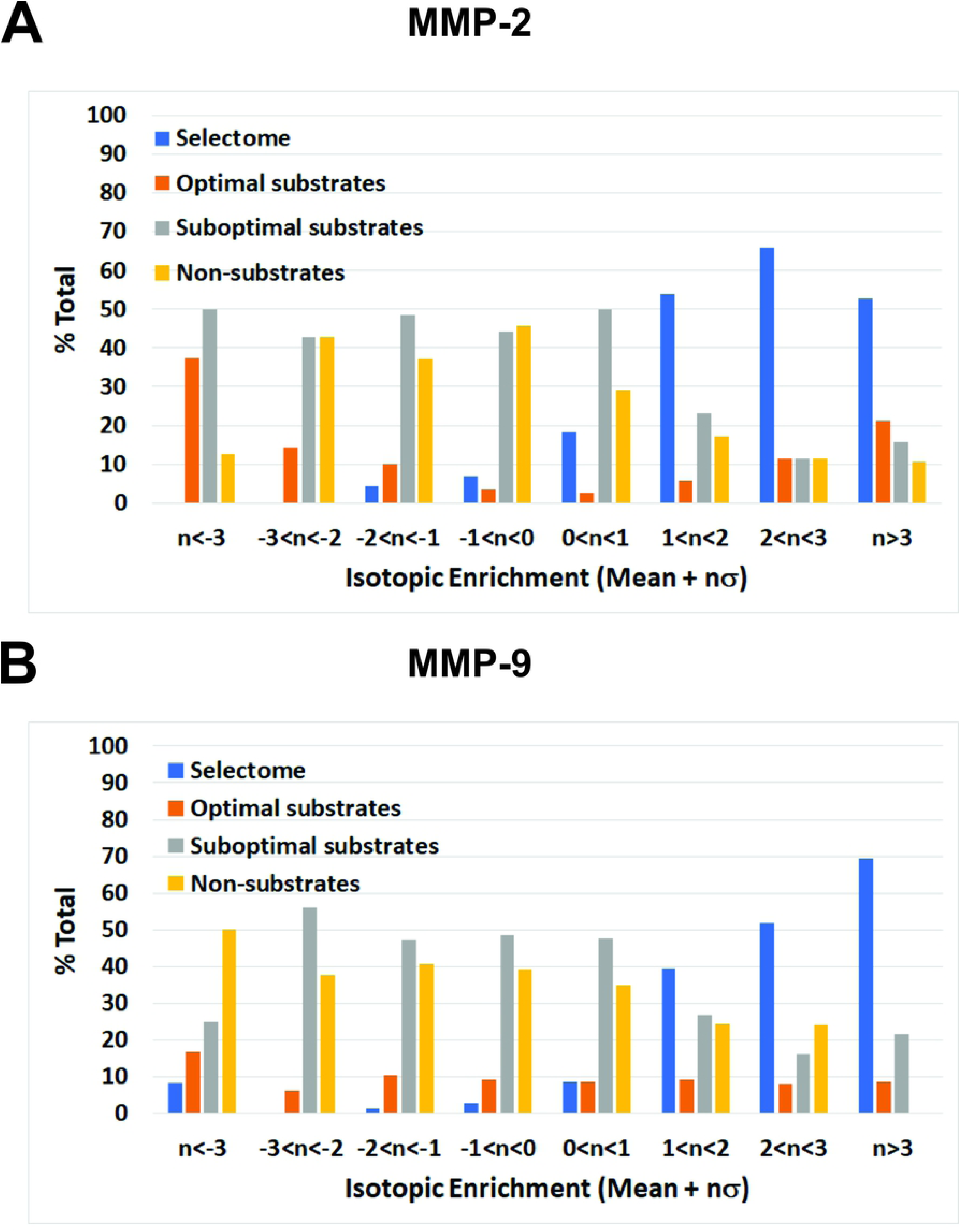
Enrichment in novel N-termini following hydrolysis by MMP-2 or 9 reflects the proportion of protein substrates in respective selectomes. Following hydrolysis with MMP-2 or 9, the secretome of HEK293 cells was labeled with TMT isobaric tags. Isotopic enrichment (IE) of the novel N-terminally labeled peptides in the MMP-treated samples relative to the untreated controls, was determined by LC/MS analysis of tryptic digests of the labeled secretomes (See text and Materials and Methods for details). N-terminally labeled peptides were split into groups based on their IE, expressed as multiples of the standard deviation away from the population mean. The percentages of N-termini corresponding to tetramer clusters with RP values defining the selectomes (≥4.5 for MMP-2 and ≥4.7 for MMP-9), optimal substrates (1<RP<4.5 for MMP-2 and 1<RP<4.7 for MMP-9), suboptimal substrates (0<RP<1 for both MMP-2 and 9) and non-substrates (RP=0) in each EI group are plotted for MMP-2 **(A)** and MMP-9 **(B)** treated secretomes.

MEROPS is a rich source of data for specificity profiling of proteases [2, 11]. It is therefore of interest to compare how the information on MMP-2 and 9 cleavages compiled from a wide variety of experimental studies is matched by our criteria for specificity. As can be seen in S17 Table, 65 and 50% of all cleavages are optimal substrates (RP>1) out of which 55% and 36% belong to the selectomes of MMP-2 and MMP-9, respectively. Of the cleavages with RP values below 1, 23 and 32% constitute suboptimal substrates and 12 and 18% are not substrates of MMP-2 and 9 based on our criteria. Of the published “physiologic” substrates, 53 and 45% are optimal, out of which 44 and 31% belong to the selectomes of MMP-2 and MMP-9, respectively. 31 and 35% of cut sites in the “physiologic” substrate category belong to the sets of suboptimal substrates and 16 and 20% are not substrates of MMP-2 and MMP-9.

In summary, based on our analysis, the catalytic cleft specificity is an important determinant of substrate fitness, which cannot be ignored. If a cleavage site does not match the catalytic cleft specificity, then it must be a relatively poor substrate representing a bystander proteolytic event or must have a large exosite contribution to its fitness. This knowledge potentially constitutes a significant advance in understanding of the mechanistic basis of MMP biology and needs further studying. So, it is useful to know if experimentally determined cleavages can be confirmed by the selectome analysis in order to decide what category a given substrate belongs to and if further study is needed.

## Discussion

### The concept of “selectome”

We developed a concept of “selectome”, i.e. a set of substrates, which, only as a whole, represents specificity of the catalytic cleft of a protease. Since, as a rule, proteases cleave multiple substrates with varying specificity constants (k_cat_/K_M_), it would be advantageous to determine a set of substrates that can be used to uniquely and quantitatively define specificities of their catalytic clefts. To the best of our knowledge, we are the first to propose such a concept. A widely accepted term “degradome” refers to the repertoire of all natural substrates cleaved by a protease [44], which may or may not be reflective of the specificity of the catalytic cleft alone. Using an information theory-based approach, we defined “selectome” as a collection of amino acid sequences with cumulative non-zero contribution to the Kullback-Leibler divergence calculated between their probability distributions in the substrate set and the set of all possible amino acid sequences of the length determined by the number and positions of the catalytic cleft selectivity determinants. In our study, we used two closely related members of the Matrix Metalloproteinase (MMP) family (MMP-2 and 9) and a close to maximally diverse library of hexapeptides displayed on gene 3 protein of M13 phage to experimentally substantiate the concept of selectome. Since the enzymes of this family have two major selectivity determinants at S3 and S1’, together with S2 and S1 between them they form a tetramer binding unit (Fig. 1). Based on that, MMPs can theoretically have between 1 and 160,000 tetramer substrates depending on the degree of specificity they possess. We found that selectomes of MMP-2 and 9 contain 7,921 and 6,094 tetramer substrates, respectively. It is important to emphasize that the selectomes of MMP-2 and 9 constitute about 10% of all the tetramer clusters identified in the substrate sets (78,757 and 76,696, respectively) indicating that the majority of the tetramer sequences cleaved by these enzymes are relatively poor substrates and contribute little, if at all, to their specificities. Currently, one of the major roadblocks to understanding protease function lies in lack of sensitive approaches to distinguishing specificities of closely related proteases from the same families [12]. Comparative analysis of selectomes of closely related proteases in combination with structural data and prior knowledge of the composition of selectivity determinants, allows to identify the SDPs responsible for the observed differences in specificity as shown in Fig. 6. This is a very valuable aspect of the selectome-based substrate specificity profiling as it provides basis for distinguishing specificities of closely related members of protease families in order to develop selective activity probes and inhibitors.

### Quantitation of substrate specificity

We developed methodology for quantification of substrate specificity based on both the number of substrates and their relative catalytic efficiencies. We introduced a probability-based quantitative metric of the specificity constant (k_cat_/K_M_) called Relative Probability (RP). It is defined as a ratio of probabilities of finding a particular amino acid sequence in the substrate and the random sets of amino acid sequences of a given length (equation 3). The area under the curve for the distribution of the number of sequences across the entire RP/RP_Max_ range (Total Substrate Fitness or TSF) can be used as a quantitative measure of specificity of a given protease and can range from 1 for a uniquely specific MMP to 160,000 for a non-specific MMP.

Studies by another group also used information theory to quantify protease specificity but purely by evaluating frequencies of occurrence of amino acid residues at different positions relative to the scissile bond [11, 30]. Based on their analysis of the MEROPS database of protease substrates, specificities of most proteases are very broad and, in fact, close to the theoretical maximum. For instance, their take on the overall specificity of MMP-2 and 9 indicates that both are broadly specific, with Shannon entropies of 7.386 and 7.078 out of the theoretically maximal 8.0 for eight sub pockets covering S4 – S4’. Shannon entropy was calculated using log_20_ so that a non-specific binding pocket accepting all residues would have a value of 1 and a totally specific binding pocket accepting a single residue would have a value of 0. Each sub pocket’s Shannon entropies are added together to obtain the overall specificity measure.

In relative terms, based on their analysis, specificities of MMP-2 and 9 are 20^8^/20^7.386^ = 6.29-fold and 20^8^/20^7.078^ = 15.83-fold narrower than that of a protease with no specificity. Our data based on the Shannon entropy values of tetramer cluster distributions in the MMP-2 and 9 substrate sets show their specificities are 2^17.288^/2^13.93^ = 10.25-fold and 2^17.288^/2^13.67^ = 12.28-fold narrower than that of a randomly specific protease. The two analyses are in a reasonably good agreement on the overall specificities of the two enzymes. These numbers imply that approximately 10% of all peptide bonds in proteins available for cleavage are substrates of MMP-2 and 9, which makes every protein in the human proteome a potential target with at least one cleavage site. Based on the TSF quantitation of selectomes, MMP-2 and 9 are 160,000/691 = 232 and 160,000/511= 313-fold more specific than a non-specific protease, respectively. This implies that if one knows the composition of the selectome of a protease and takes into account the substrate numbers and relative fitness to quantify specificity, both enzymes are much more specific than can be assessed just based on residue frequencies relative to the scissile bond of all the cleavage sites derived from the substrate sequences.

### Relevance to proteolysis of folded proteins

Using the results of a published rigorous study [42]and our own data, we determined the relevance of our selectome-based approach to identification of cleavage sites in folded proteins. Our analysis of the published results shows that from 70 to 80% of the protein substrates identified with high confidence belong to the selectomes of MMP-2 and 9. Our own analysis of cleavages in folded proteins based on enrichment of novel N-termini following MMP treatment, demonstrates that RP above the selectome threshold for the matching P3-P1’ tetramers is the best predictor for enrichment of the corresponding N-termini greater than σ = 1 above the population mean (S16 Table).

Selectome-based quantification of substrate specificity shows that MMP-2 and 9 are ∼200-300-fold more specific than a randomly specific protease, which implies that 1 out of 200-300 peptide bonds in proteins has the same probability of being cleaved by MMP-2 and 9 as by a non-specific protease. Given the average length of a eukaryotic protein (472 residues [45]), about 2 peptide bonds per protein are potentially relevant cleavage sites of either MMP. Given that in order to be accessible to proteolysis a peptide bond has to be solvent-accessible [46] and the average solvent exposure observed in folded proteins is 20-30% [47], less than 1 peptide bond per average-size eukaryotic protein is potentially a cleavage site that belongs to the selectome of either MMP-2 or 9. This is consistent with the notion of substrate-protease co-evolution necessary for regulatory proteolysis to take place in a living organism [48], where protease activity is strictly limited to physiologically compatible levels by natural inhibition and pro-enzyme latency. A corollary to that is that the relatively poor substrates, constituting most of the tetramer clusters of MMP-2 and 9, have little relevance on the time scale of targeted proteolysis occurring under normal physiologic conditions, but might become quite important in pathological states in which protease activity is deregulated. Our analysis of the MEROPS database of MMP-2 and 9 substrates shows that a significant proportion of reported cleavages (∼50%) represent suboptimal substrates or non-substrates based on the RP classification. They can belong to off-target proteolytic events or artifacts, or possibly constitute exosite driven proteolysis by MMP-2 and 9. This information is very useful for follow-up studies to determine if exosite participation is a significant component of substrate recognition by these and other MMPs [49]. The fact that ∼80% of the P3-P1’ tetramers in the substrate sets of MMP-2 and 9 are suboptimal substrates indicates that simple accounting for amino acid diversity at individual positions of substrates is not an accurate estimate of physiologically relevant specificity. Thus, our selectome-based analysis of cleavage events in folded proteins establishes the importance of catalytic cleft specificity in protein substrate recognition by MMP-2 and 9.

### Conclusion

Work presented here establishes a new approach to studying substrate specificity of proteases and possibly other enzymes involved in posttranslational modification of proteins [50]. It is based on statistically saturated data sets and a new way of applying information theory to quantitatively define substrate specificity of proteases by employing a novel concept of “selectome”. In practical terms, this approach can be invaluable for developing highly selective activity probes and inhibitors for closely related members of large protease families. By providing a measure of catalytic efficiency, our approach can also be used to help determine which cleavages in folded proteins represent physiologic and pathologic targets and which are bystander proteolytic events.

## Materials and Methods

### Expression and Purification of Recombinant Catalytic Domains and activity assays

The recombinant catalytic domains of MMP-2 and -9 were expressed in HEK293 cells stably transfected with respective constructs and purified from serum-free culture medium using Gelatin Sepharose 4B (GE Healthcare) as described in [35, 39]. Following activation with APMA (Sigma-Aldrich), the amount of active enzyme was determined by active site titration using GM6001 (Sigma-Aldrich) [35]. The k_cat_/K_M_ values of 100 peptides derived from phage substrates (S7, Table) were determined in triplicate as described in [33].

### Substrate phage selections and NGS analysis

Selection of substrate phage was performed as described in [10]. Briefly, 5 × 10^11^ phage particles were mixed with a protease at 200 nM of active enzyme in 0.5 mL of the reaction buffer and incubated for 2 h at 37 °C. A reaction without addition of protease was performed as a control. Following incubation, MMP activity was halted by addition of GM6001. Phage with uncleaved FLAG-tag were removed by immunodepletion using M2 anti-FLAG monoclonal antibody (Sigma-Aldrich) coupled to epoxy-activated M-450 magnetic beads (Invitrogen). The extent of immunodepletion in the protease treated and untreated control samples was determined by ELISA. Unbound phage were propagated, purified by PEG precipitation and used in the second round of selection. Phage ssDNA from the control and protease selections was purified from 8 × 10^12^ phage particles using phenol-chloroform extraction. The region harboring the randomized hexamer and flanking constant tags was amplified from 1 µg ssDNA for 5 cycles using the Q5 Hot Start High-Fidelity PCR Master Mix (New England Biolabs) and the following primers: 5’-ACGACGACGACAAACCCG-3’ forward and 5’-AACAGTTTCGGCCCCAGA-3’ reverse. The NGS library was prepared using NEBNext Ultra II DNA Library Prep Kit for Illumina and NEBNext Multiplex Oligos for Illumina. Amplicons were cleaned up using AMPure XP beads (Beckman Coulter) and subjected to NGS analysis performed at the Genomics Core of the Sanford-Burnham-Prebys Medical Discovery Institute. In total, 324,285,851 reads were generated to sequence the naïve phage display library, and 27,221, 954 and 34,786,206 to sequence the MMP-2 and MMP-9 substrate selections, respectively. We developed a series of Linux shell scripts and Fortran programs for processing and analysis of NGS data. FASTQC program (Andrews, S. (2010). FastQC: AQuality Control Tool for High Throughput Sequence Data [Online], available online at: http://www.bioinformatics.babraham.ac.uk/projects/fastqc/) was used to check for quality of sequencing data. No low-quality sequences were detected in the NGS raw fastq files. The DNA sequences were translated in forward and reverse directions using all three reading frames. Sequences of the variable hexamer region were accepted only if they were flanked by the constant tag flanking sequences. Sequences belonging to the variable hexamer region were used for all downstream analyses. Sequences of peptide hexamers found in the untreated controls were removed from the downstream analysis of substrate selections. These analyses include sorting, assigning hexamer sequences to tetramer clusters, calculating probabilities of tetramers, deriving Shannon entropy and Kullback-Leibler divergence.

### Proteomic identification of novel N-termini in folded proteins

#### Sample preparation

HEK293 cells were seeded in 150-mm tissue culture dishes (Corning) and grown to confluence. The confluent cultures were washed with PBS several times to remove serum proteins and kept in serum-free DMEM supplemented with 1 µM GM6001 for 72 hours at 37 C, 5% CO_2_. The conditioned media was centrifuged at 10,000 x g for 30 minutes at 4 C and concentrated **∼**100-fold using Centricon Plus-70 (AMD Millipore) 3-kDa m.w. cut-off ultrafilter, and the buffer was exchanged to 50 mM HEPES pH 7.0 containing 150 mM NaCl and 10 mM CaCl_2_ (Reaction Buffer) using a PD-10 desalting column (GE Healthcare). Aliquots containing 240 µg total protein were incubated at 37 C in the presence of 750 nM MMP2 or 9 or the Reaction Buffer alone for 2 hours. Each reaction was carried out in duplicate. The reactions were stopped by addition of 20 mM EDTA and subjected to denaturation/reduction (4M Urea and 10 mM TCEP, 1 h, 22°C) and alkylation (20 mM iodoacetamide, 30 min, 22°C, in dark). Each sample was labeled for 2 h at 22°C with a different TMT-tag obtained from a TMT10plex kit (Thermo Fisher Scientific), and then quenched (0.27% hydroxylamine, 15 min, 22°C). The buffer control samples were labeled with TMT-127N and TMT-127C, the MMP-2 treated samples were labeled with TMT-128N and TMT-128C and the MMP-9 treated sampled were labeled with TMT-129C and TMT-130C. The TMT-labeled samples were pooled together, and subjected to TCA precipitation (20% TCA, 16 h, 4°C). Each pellet was washed with -20°C acetone, recovered in 50 mM Tris-HCl pH 8.0, and then digested with trypsin/Lys-C (Promega) for 16 h at 37°C. Digestions were stopped with 0.2% TFA and centrifuged to remove insoluble material. The supernatants were then desalted using Sep-Pak C18 cartridge (50 mg), and then lyophilized.

#### Offline fractionation

Dried pooled sample was reconstituted in 20 mM ammonium formate pH ∼10 and fractionated using a Waters Acquity BEH C18 column (2.1x 15 cm, 1.7 µm pore size) mounted on an M-Class Ultra Performance Liquid Chromatography (UPLC) system (Waters). Peptides were then separated using a 35-min gradient: 5% to 18% B in 3 min, 18% to 36% B in 20 min, 36% to 46% B in 2 min, 46% to 60% B in 5 min, and 60% to 70% B in 5 min (A=20 mM ammonium formate, pH 10; B = 100% ACN). A total of 32 fractions were collected and pooled in a non-contiguous manner into 16 total fractions. Pooled fractions were dried to completeness in a SpeedVac concentrator prior to mass spectrometry analysis

#### LC-MS/MS analysis

Dried peptide fractions were reconstituted with 2% ACN-0.1% FA and analyzed by LC-MS/MS using a Proxeon EASY nanoLC system (Thermo Fisher Scientific) coupled to a Q-Exactive Plus mass spectrometer (Thermo Fisher Scientific). Peptides were separated using an analytical C18 Acclaim PepMap column (75µm x 250 mm, 2µm particles; Thermo Scientific) at a flow rate of 300 µl/min using a 58-min gradient: 1% to 6% B in 1 min, 6% to 23% B in 35 min, and 23% to 34% B in 22 min (A= FA, 0.1%; B=80% ACN: 0.1% FA). The mass spectrometer was operated in positive data-dependent acquisition mode. MS1 spectra were measured with a resolution of 70,000 (AGC target: 1e6; mass range: 350-1700 m/z). Up to 12 MS2 spectra per duty cycle were triggered, fragmented by HCD, and acquired with a resolution of 17,500 (AGC target 1e5, isolation window; 1.2 m/z; normalized collision: 32) Dynamic exclusion was enabled with a duration of 25 sec.

#### Analysis of the proteomics data

Raw files were analyzed using MaxQuant software version 1.6.8.0 and MS/MS spectra were searched against the *Homo sapiens* Uniprot protein sequence database (downloaded in January 2019). The false discovery rate (FDR) filter for spectrum and protein identification was set to 1%. Carbamidomethylation of cysteines was searched as a fixed modification, while oxidation of methionines and N-terminal acetylation were searched as variable modifications. Enzyme was set to trypsin in a semispecific mode and a maximum of two missed cleavages was allowed for searching. To obtain quantification of the TMT-labeled N-termini, two independent searches were performed – one with TMT10plex MS2 N-terminal reporter quantification and the other for quantification of N-terminal and lysine sidechain. The results were combined to obtain the complete set of the N-terminally labeled peptides

## Supporting information

**S1 Fig. Most hexamers in substrate selections of MMP-2 and 9 belong to few tetramer clusters**.

Tetramer clusters in substrate selections of MMP-2 and 9 were grouped into 10% bins based on their RP values relative to the maximum. The numbers of tetramer clusters in each bin (**A**) and the numbers of hexamers in the corresponding tetramer clusters (**B**) are plotted as a function of the RP interval they belong to.

**S2 Fig. Specificity profiles based on the unique and overlapping tetramer clusters of MMP-2 and 9 selectomes as a function of RP/RP**_**Max**_. Logo plots demonstrate the composition of substrates across P5 to P3’ positions as a function of RP/RP_Max_. Peptide hexamers belonging to tetramer clusters in the selectomes of MMP-2 and 9 were aligned across P3-P1’ positions and divided into 10 groups based on their RP values relative to the maximum (RP/RP_Max_). First and second columns of logo plots correspond to unique substrates of MMP-2 and 9 selectomes, respectively. The third and fourth columns of logo plots represent the common set of MMP-2 and 9 selectomes, respectively. The RP values for the corresponding tetramer clusters have been calculated either according to MMP-2 (MMP-2&9/2) or MMP-9 (MMP-2&9/9) ranking.

**S1 Table**. Information about the numbers of hexamers and resultant tetramer clusters in the naïve library and MMP-2 and 9 substrate selections.

**S2 Table**. List of all tetramer clusters and corresponding aligned hexamers in the naive phage display library. Tetramer amino acid sequence, rank and the number of hexamers in it are shown in the header for each tetramer cluster.

**S3 Table**. List of all tetramer clusters and corresponding hexamers aligned along P3-P1’ positions of MMP-2 substrates. Tetramer amino acid sequence, rank and the number of hexamers in it are shown in the header for each tetramer cluster.

**S4 Table**. List of all tetramer clusters and corresponding hexamers aligned along P3-P1’ positions of MMP-9 substrates. Tetramer amino acid sequence, rank and the number of hexamers in it are shown in the header for each tetramer cluster.

**S5 Table**. Statistical information about tetramer clusters for MMP-2. The table contains information about: a) the amino acid sequence of tetramer cluster, b) rank of the tetramer cluster calculated using relative probability, c) number of hexamers in a cluster from MMP set and d) number of hexamers in the corresponding cluster in naïve library, e) ratio of hexamer numbers in the MMP substrate set and the naïve library (%), f) probability of a tetramer in MMP set, g) probability of corresponding tetramer in naïve library, h) relative probability calculated as a ratio of (f) and (g) probabilities, i) individual contribution of each tetramer cluster to Kullback-Leibler (K-L) divergence, j) cumulative values of K-L divergence over tetramer clusters, k) individual contribution of each tetramer cluster to Shannon entropy, l) cumulative values of Shannon entropy over tetramer clusters. The resultant value of K-L divergence and Shannon entropy for MMP set and naïve library is provided at the end of each table.

**S6 Table**. Statistical information about tetramer clusters for MMP-9. See S5 Table for explanation of table content.

**S7 Table**. Table of 1369 peptide set derived and published previously [10] for MMP-2 and 9, for which positions of scissile bonds and K_(obs)_ values were determined experimentally. Each tab, corresponding to MMP-2 and 9, respectively, contains information about: a) the sequence of dodecamer substrate with marked cleavage position, b) the corresponding amino acid tetramer sequences for P3-P1’ positions, c) rank of tetramer based on RP value, d) RP value, e) measured K_(obs)_ (M^-1^s^-1^).

**S8 Table**. Statistical performance of tetramer approach to analysis of cleavages in set of 1369 substrates [10] for MMP-2 and MMP-9. Only nonredundant sets of peptide substrates have been selected for statistical assessment. The value of K_(obs)_ (M^-1^s^-1^) for each tetramer has been calculated as an average value over all redundant entries. The calculations have been performed for three different thresholds related to K-L divergence analysis: a) for RP above 4.5 or 4.7 for MMP-2 or 9, respectively (“selectome”), b) for RP value above 1 (set of optimal substrates), and c) for RP above 0, which includes all substrates.

**S9 Table**. Table of 100 peptide set derived from phage substrates selection for MMP-2 and 9, for which k_cat_/K_M_ values were experimentally determined. Each tab, corresponding to MMP-2 and 9, respectively, contains information about a) the sequence of hexamer substrate, b) the corresponding amino acid tetramer sequences for P3-P1’ positions, c) rank of tetramer based on RP value, d) relative probability (RP), e) measured k_cat_/K_M_ (M^-1^s^-1^), f-g) standard deviation and standard error for measured k_cat_/K_M_ (M^-1^s^-1^), based on triplicate experiments for each experiment.

**S10 Table**. Analysis of combined selectomes of MMP-2 and 9. **A**. List of unique tetramer clusters and corresponding hexamer sequences aligned across P3-P1’ positions of substrates belonging to MMP-2 selectome. Tetramers are ranked according to MMP-2 relative probability. The number of hexamers in a tetramer cluster depends on which MMP ranking was applied. Information about the tetramer amino acids sequence, rank, number of hexamers and relative probability of a cluster is provided in the header for each tetramer cluster.

**S11 Table**. Analysis of combined selectomes of MMP-2 and 9. List of unique tetramer clusters and corresponding hexamer sequences aligned across P3-P1’ positions of substrates belonging to MMP-9 selectome. Tetramers are ranked according to MMP-9 relative probability. Information about the tetramer amino acids sequence, rank, number of hexamers and relative probability of a cluster is provided in the header for each tetramer cluster.

**S12 Table**. Analysis of combined selectomes of MMP-2 and 9. List of tetramer clusters common between the selectomes of MMP-2 and 9 together with the corresponding hexamer sequences aligned across P3-P1’ positions of substrates. Tetramers are ranked according to MMP-2 relative probability. Information about the tetramer amino acids sequence, rank, number of hexamers and relative probability of a cluster is provided in the header of each tetramer cluster.

**S13 Table**. Analysis of combined selectomes of MMP-2 and 9. List of tetramer clusters common between the selectomes of MMP-2 and 9 together with the corresponding hexamer sequences aligned across P3-P1’ positions of substrates. Tetramers are ranked according to MMP-9 relative probability. Information about the tetramer amino acids sequence, rank, number of hexamers and relative probability of a cluster is provided in the header of each tetramer cluster.

**S14 Table**. Identification of cleavage sites using tetramer projection in MMP-2 (**A**) and MMP-9 (**B**) substrates, determined by Prudova *et al*. (2010) [42]. Each table contains information about tetramer sequence projected onto the cleavage site, its rank and relative probability.

**S15 Table**. Tetramer matching to novel N-termini in proteins secreted by HEK293 cells following treatment with MMP-2 (**A**) and MMP-9 (**B**). Each table contains sequences of deduced cleavages grouped according to the ranges of isotopic enrichment of the corresponding N-terminally labeled peptides and relative probability values of the matching tetramers. For each identified cleavage the following information has been provided: the rank and RP of the matching tetramer, isotopic enrichment (log_2_(ratio)) and p-value for each N-terminally labeled peptide based on duplicate determinations and respective protein ids.

**S16 Table**. Binary classification of cleavage sites in HEK293 cell secretomes following treatment with MMP-2 (A) and MMP-9 (B). Cleavages detected in HEK293 cell secretome have been grouped based on the value of an RP threshold (a) 4.5 (MMP-2) or 4.7 (MMP-9), based on K-L defined selectomes, b) 1, for optimal substrates, and c) all non-zero – all substrates, including the suboptimal) and n x σ distance away from the average value of log_2_ IE (isotopic enrichment). For each group the following binary classifiers have been used: TP - number of cases for which log_2_ IE is above a certain n x σ, and RP of associated tetramers is above a specified threshold; TN - number of cases for which log_2_ IE is below the n x σ, and RP is below the threshold; FN - number of cases for which log_2_ IE is above the n x σ, and RP is below the threshold; FP - number of cases for which log_2_ IE is below the n x σ, and RP is above the threshold. For each group, sensitivity, specificity, accuracy, FP rate, precision and Matthews correlation coefficient (MCC) have been determined.

**S17 Table**. Tetramer annotation of cleavage sites in MMP-2 and 9 substrates collected from the MEROPS database. The data have been divided into two groups – those annotated as physiologic substrates (**A**) – MMP2, (**C)** – MMP9, and all substrates (**B**) – MMP2, (**D**) – MMP9. Each table contains information about octamer sequences covering P4-P4’ positions of substrates reported in MEROPS, the corresponding tetramers, their ranks and relative probabilities, as well as MEROPS annotation relating the cleavage site to its position in a protein or analyzed polypeptide.

## Acknowledgements

We would like to thank the Proteomics Facility at the Sanford-Burnham-Prebys Medical Discovery Institute and its director Dr. Alex Rosa Campos for the proteomic analysis. We also would like to thank the Genomics Core Facility at the Sanford-Burnham-Prebys Medical Discovery Institute and its director Dr. Brian James for the NGS sequencing support. We would like to thank Dr. Marat Kazanov of The Research and Training Center on Bioinformatics, Institute for Information Transmission Problems, Russian Academy of Sciences, Moscow 127994 for helpful comments and advice.

## References

1. Oda K. New families of carboxyl peptidases: serine-carboxyl peptidases and glutamic peptidases. J Biochem. 2012;151(1):13-25. Epub 2011/10/22. doi: 10.1093/jb/mvr129. PubMed PMID: 22016395.

2. Rawlings ND, Barrett AJ, Thomas PD, Huang X, Bateman A, Finn RD. The MEROPS database of proteolytic enzymes, their substrates and inhibitors in 2017 and a comparison with peptidases in the PANTHER database. Nucleic Acids Res. 2018;46(D1):D624-D32. Epub 2017/11/18. doi: 10.1093/nar/gkx1134. PubMed PMID: 29145643; PubMed Central PMCID: PMCPMC5753285.

3. Doucet A, Butler GS, Rodriguez D, Prudova A, Overall CM. Metadegradomics: toward in vivo quantitative degradomics of proteolytic post-translational modifications of the cancer proteome. Mol Cell Proteomics. 2008;7(10):1925-51. Epub 2008/07/04. doi: 10.1074/mcp.R800012-MCP200. PubMed PMID: 18596063.

4. Vizovisek M, Vidmar R, Drag M, Fonovic M, Salvesen GS, Turk B. Protease Specificity: Towards In Vivo Imaging Applications and Biomarker Discovery. Trends Biochem Sci. 2018;43(10):829-44. Epub 2018/08/12. doi: 10.1016/j.tibs.2018.07.003. PubMed PMID: 30097385.

5. Green D. Coagulation cascade. Hemodial Int. 2006;10 Suppl 2:S2-4. Epub 2006/10/07. doi: 10.1111/j.1542-4758.2006.00119.x. PubMed PMID: 17022746.

6. Lopez-Otin C, Hunter T. The regulatory crosstalk between kinases and proteases in cancer. Nat Rev Cancer. 2010;10(4):278-92. Epub 2010/03/20. doi: 10.1038/nrc2823. PubMed PMID: 20300104.

7. Budenholzer L, Cheng CL, Li Y, Hochstrasser M. Proteasome Structure and Assembly. J Mol Biol. 2017;429(22):3500-24. Epub 2017/06/07. doi: 10.1016/j.jmb.2017.05.027. PubMed PMID: 28583440; PubMed Central PMCID: PMCPMC5675778.

8. Dix MM, Simon GM, Wang C, Okerberg E, Patricelli MP, Cravatt BF. Functional interplay between caspase cleavage and phosphorylation sculpts the apoptotic proteome. Cell. 2012;150(2):426-40. Epub 2012/07/24. doi: 10.1016/j.cell.2012.05.040. PubMed PMID: 22817901; PubMed Central PMCID: PMCPMC3569040.

9. Corral J, Vicente V, Carrell RW. Thrombosis as a conformational disease. Haematologica. 2005;90(2):238-46. Epub 2005/02/16. PubMed PMID: 15710578.

10. Ratnikov BI, Cieplak P, Gramatikoff K, Pierce J, Eroshkin A, Igarashi Y, et al. Basis for substrate recognition and distinction by matrix metalloproteinases. Proc Natl Acad Sci U S A. 2014;111(40):E4148-55. Epub 2014/09/24. doi: 10.1073/pnas.1406134111. PubMed PMID: 25246591; PubMed Central PMCID: PMCPMC4210027.

11. Fuchs JE, von Grafenstein S, Huber RG, Margreiter MA, Spitzer GM, Wallnoefer HG, et al. Cleavage entropy as quantitative measure of protease specificity. PLoS Comput Biol. 2013;9(4):e1003007. Epub 2013/05/03. doi: 10.1371/journal.pcbi.1003007. PubMed PMID: 23637583; PubMed Central PMCID: PMCPMC3630115.

12. Kasperkiewicz P, Poreba M, Groborz K, Drag M. Emerging challenges in the design of selective substrates, inhibitors and activity-based probes for indistinguishable proteases. FEBS J. 2017;284(10):1518-39. Epub 2017/01/05. doi: 10.1111/febs.14001. PubMed PMID: 28052575; PubMed Central PMCID: PMCPMC7164106.

13. Klein T, Eckhard U, Dufour A, Solis N, Overall CM. Proteolytic Cleavage-Mechanisms, Function, and “Omic” Approaches for a Near-Ubiquitous Posttranslational Modification. Chem Rev. 2018;118(3):1137-68. Epub 2017/12/22. doi: 10.1021/acs.chemrev.7b00120. PubMed PMID: 29265812.

14. Drag M, Salvesen GS. Emerging principles in protease-based drug discovery. Nat Rev Drug Discov. 2010;9(9):690-701. Epub 2010/09/03. doi: 10.1038/nrd3053. PubMed PMID: 20811381; PubMed Central PMCID: PMCPMC2974563.

15. Poreba M, Szalek A, Kasperkiewicz P, Rut W, Salvesen GS, Drag M. Small Molecule Active Site Directed Tools for Studying Human Caspases. Chem Rev. 2015;115(22):12546-629. Epub 2015/11/10. doi: 10.1021/acs.chemrev.5b00434. PubMed PMID: 26551511; PubMed Central PMCID: PMCPMC5610424.

16. auf dem Keller U, Doucet A, Overall CM. Protease research in the era of systems biology. Biol Chem. 2007;388(11):1159-62. Epub 2007/11/03. doi: 10.1515/BC.2007.146. PubMed PMID: 17976008.

17. Diamond SL. Methods for mapping protease specificity. Curr Opin Chem Biol. 2007;11(1):46-51. Epub 2006/12/13. doi: 10.1016/j.cbpa.2006.11.021. PubMed PMID: 17157549.

18. Turk BE, Huang LL, Piro ET, Cantley LC. Determination of protease cleavage site motifs using mixture-based oriented peptide libraries. Nat Biotechnol. 2001;19(7):661-7. Epub 2001/07/04. doi: 10.1038/90273. PubMed PMID: 11433279.

19. Chen S, Yim JJ, Bogyo M. Synthetic and biological approaches to map substrate specificities of proteases. Biol Chem. 2019;401(1):165-82. Epub 2019/10/23. doi: 10.1515/hsz-2019-0332. PubMed PMID: 31639098.

20. Poreba M, Salvesen GS, Drag M. Synthesis of a HyCoSuL peptide substrate library to dissect protease substrate specificity. Nat Protoc. 2017;12(10):2189-214. Epub 2017/09/22. doi: 10.1038/nprot.2017.091. PubMed PMID: 28933778.

21. Matthews DJ, Wells JA. Substrate phage: selection of protease substrates by monovalent phage display. Science. 1993;260(5111):1113-7. Epub 1993/05/21. doi: 10.1126/science.8493554. PubMed PMID: 8493554.

22. Schilling O, Overall CM. Proteome-derived, database-searchable peptide libraries for identifying protease cleavage sites. Nat Biotechnol. 2008;26(6):685-94. Epub 2008/05/27. doi: 10.1038/nbt1408. PubMed PMID: 18500335.

23. Kretz CA, Dai M, Soylemez O, Yee A, Desch KC, Siemieniak D, et al. Massively parallel enzyme kinetics reveals the substrate recognition landscape of the metalloprotease ADAMTS13. Proc Natl Acad Sci U S A. 2015;112(30):9328-33. Epub 2015/07/15. doi: 10.1073/pnas.1511328112. PubMed PMID: 26170332; PubMed Central PMCID: PMCPMC4522773.

24. Kretz CA, Tomberg K, Van Esbroeck A, Yee A, Ginsburg D. High throughput protease profiling comprehensively defines active site specificity for thrombin and ADAMTS13. Sci Rep. 2018;8(1):2788. Epub 2018/02/13. doi: 10.1038/s41598-018-21021-9. PubMed PMID: 29434246; PubMed Central PMCID: PMCPMC5809430.

25. Rentero Rebollo I, Sabisz M, Baeriswyl V, Heinis C. Identification of target-binding peptide motifs by high-throughput sequencing of phage-selected peptides. Nucleic Acids Res. 2014;42(22):e169. Epub 2014/10/29. doi: 10.1093/nar/gku940. PubMed PMID: 25348396; PubMed Central PMCID: PMCPMC4267670.

26. Cornish-Bowden A. Enzyme specificity: its meaning in the general case. J Theor Biol. 1984;108(3):451-7. Epub 1984/06/07. doi: 10.1016/s0022-5193(84)80045-4. PubMed PMID: 6748701.

27. Lehninger AL, Cox, Michael M.Nelson, David L. Lehninger principles of biochemistry. New York: W.H. Freeman; 2008.

28. Schilling O, auf dem Keller U, Overall CM. Protease specificity profiling by tandem mass spectrometry using proteome-derived peptide libraries. Methods Mol Biol. 2011;753:257-72. Epub 2011/05/24. doi: 10.1007/978-1-61779-148-2_17. PubMed PMID: 21604128.

29. Eckhard U, Huesgen PF, Schilling O, Bellac CL, Butler GS, Cox JH, et al. Active site specificity profiling of the matrix metalloproteinase family: Proteomic identification of 4300 cleavage sites by nine MMPs explored with structural and synthetic peptide cleavage analyses. Matrix Biol. 2016;49:37-60. Epub 2015/09/27. doi: 10.1016/j.matbio.2015.09.003. PubMed PMID: 26407638.

30. Schauperl M, Fuchs JE, Waldner BJ, Huber RG, Kramer C, Liedl KR. Characterizing Protease Specificity: How Many Substrates Do We Need? PLoS One. 2015;10(11):e0142658. Epub 2015/11/13. doi: 10.1371/journal.pone.0142658. PubMed PMID: 26559682; PubMed Central PMCID: PMCPMC4641643.

31. Schechter I, Berger A. On the size of the active site in proteases. I. Papain. 1967. Biochem Biophys Res Commun. 2012;425(3):497-502. Epub 2012/08/29. doi: 10.1016/j.bbrc.2012.08.015. PubMed PMID: 22925665.

32. Maskos K. Crystal structures of MMPs in complex with physiological and pharmacological inhibitors. Biochimie. 2005;87(3-4):249-63. Epub 2005/03/23. doi: 10.1016/j.biochi.2004.11.019. PubMed PMID: 15781312.

33. Chen EI, Kridel SJ, Howard EW, Li W, Godzik A, Smith JW. A unique substrate recognition profile for matrix metalloproteinase-2. J Biol Chem. 2002;277(6):4485-91. Epub 2001/11/06. doi: 10.1074/jbc.M109469200. PubMed PMID: 11694539.

34. Kridel SJ, Sawai H, Ratnikov BI, Chen EI, Li W, Godzik A, et al. A unique substrate binding mode discriminates membrane type-1 matrix metalloproteinase from other matrix metalloproteinases. J Biol Chem. 2002;277(26):23788-93. Epub 2002/04/18. doi: 10.1074/jbc.M111574200. PubMed PMID: 11959855.

35. Kridel SJ, Chen E, Kotra LP, Howard EW, Mobashery S, Smith JW. Substrate hydrolysis by matrix metalloproteinase-9. J Biol Chem. 2001;276(23):20572-8. Epub 2001/03/30. doi: 10.1074/jbc.M100900200. PubMed PMID: 11279151.

36. S. Kullback RAL. On information and sufficiency.. Ann Math Statist. 1951;55 79–86.

37. Crooks GE, Hon G, Chandonia JM, Brenner SE. WebLogo: a sequence logo generator. Genome Res. 2004;14(6):1188-90. Epub 2004/06/03. doi: 10.1101/gr.849004. PubMed PMID: 15173120; PubMed Central PMCID: PMCPMC419797.

38. Eckhard U, Huesgen PF, Schilling O, Bellac CL, Butler GS, Cox JH, et al. Active site specificity profiling datasets of matrix metalloproteinases (MMPs) 1, 2, 3, 7, 8, 9, 12, 13 and 14. Data Brief. 2016;7:299-310. Epub 2016/03/17. doi: 10.1016/j.dib.2016.02.036. PubMed PMID: 26981551; PubMed Central PMCID: PMCPMC4777984.

39. Chen EI, Li W, Godzik A, Howard EW, Smith JW. A residue in the S2 subsite controls substrate selectivity of matrix metalloproteinase-2 and matrix metalloproteinase-9. J Biol Chem. 2003;278(19):17158-63. Epub 2003/02/20. doi: 10.1074/jbc.M210324200. PubMed PMID: 12591933.

40. Nishimura H. Renin-angiotensin system in vertebrates: phylogenetic view of structure and function. Anat Sci Int. 2017;92(2):215-47. Epub 2016/10/09. doi: 10.1007/s12565-016-0372-8. PubMed PMID: 27718210.

41. Kawaguchi M, Inoue K, Iuchi I, Nishida M, Yasumasu S. Molecular co-evolution of a protease and its substrate elucidated by analysis of the activity of predicted ancestral hatching enzyme. BMC Evol Biol. 2013;13:231. Epub 2013/10/29. doi: 10.1186/1471-2148-13-231. PubMed PMID: 24161109; PubMed Central PMCID: PMCPMC3819744.

42. Prudova A, auf dem Keller U, Butler GS, Overall CM. Multiplex N-terminome analysis of MMP-2 and MMP-9 substrate degradomes by iTRAQ-TAILS quantitative proteomics. Mol Cell Proteomics. 2010;9(5):894-911. Epub 2010/03/23. doi: 10.1074/mcp.M000050-MCP201. PubMed PMID: 20305284; PubMed Central PMCID: PMCPMC2871422.

43. Graham FL, Smiley J, Russell WC, Nairn R. Characteristics of a human cell line transformed by DNA from human adenovirus type 5. J Gen Virol. 1977;36(1):59-74. Epub 1977/07/01. doi: 10.1099/0022-1317-36-1-59. PubMed PMID: 886304.

44. Lopez-Otin C, Overall CM. Protease degradomics: a new challenge for proteomics. Nat Rev Mol Cell Biol. 2002;3(7):509-19. Epub 2002/07/03. doi: 10.1038/nrm858. PubMed PMID: 12094217.

45. Tiessen A, Perez-Rodriguez P, Delaye-Arredondo LJ. Mathematical modeling and comparison of protein size distribution in different plant, animal, fungal and microbial species reveals a negative correlation between protein size and protein number, thus providing insight into the evolution of proteomes. BMC Res Notes. 2012;5:85. Epub 2012/02/03. doi: 10.1186/1756-0500-5-85. PubMed PMID: 22296664; PubMed Central PMCID: PMCPMC3296660.

46. Kazanov MD, Igarashi Y, Eroshkin AM, Cieplak P, Ratnikov B, Zhang Y, et al. Structural determinants of limited proteolysis. J Proteome Res. 2011;10(8):3642-51. Epub 2011/06/21. doi: 10.1021/pr200271w. PubMed PMID: 21682278; PubMed Central PMCID: PMCPMC3164237.

47. Sprang S, Yang D, Fletterick RJ. Solvent accessibility properties of complex proteins. Nature. 1979;280(5720):333-5. Epub 1979/07/26. doi: 10.1038/280333a0. PubMed PMID: 460408.

48. Timmer JC, Zhu W, Pop C, Regan T, Snipas SJ, Eroshkin AM, et al. Structural and kinetic determinants of protease substrates. Nat Struct Mol Biol. 2009;16(10):1101-8. Epub 2009/09/22. doi: 10.1038/nsmb.1668. PubMed PMID: 19767749; PubMed Central PMCID: PMCPMC4042863.

49. Van Doren SR. Matrix metalloproteinase interactions with collagen and elastin. Matrix Biol. 2015;44-46:224-31. Epub 2015/01/21. doi: 10.1016/j.matbio.2015.01.005. PubMed PMID: 25599938; PubMed Central PMCID: PMCPMC4466143.

50. Ivry SL, Meyer NO, Winter MB, Bohn MF, Knudsen GM, O’Donoghue AJ, et al. Global substrate specificity profiling of post-translational modifying enzymes. Protein Sci. 2018;27(3):584-94. Epub 2017/11/24. doi: 10.1002/pro.3352. PubMed PMID: 29168252; PubMed Central PMCID: PMCPMC5818756.

